# Pseudouridine-dependent ribosome biogenesis regulates translation of polyglutamine proteins during *Drosophila* oogenesis

**DOI:** 10.1101/2022.07.07.499147

**Authors:** Shane Breznak, Yingshi Peng, Limin Deng, Noor M. Kotb, Zachary Flamholz, Ian T. Rapisarda, Elliot T. Martin, Kara A. LaBarge, Dan Fabris, Elizabeth R. Gavis, Prashanth Rangan

## Abstract

Stem cells in many systems, including *Drosophila* germline stem cells (GSCs), increase ribosome biogenesis and translation during terminal differentiation. Here, we show that pseudouridylation of ribosomal RNA (rRNA) mediated by the H/ACA box is required for ribosome biogenesis and oocyte specification. Reducing ribosome levels during differentiation decreased the translation of a subset of mRNAs that are enriched for CAG repeats and encode polyglutamine-containing proteins, including differentiation factors such as RNA-binding Fox protein 1. Moreover, ribosomes were enriched at CAG repeats within transcripts during oogenesis. Increasing TOR activity to elevate ribosome levels in H/ACA box-depleted germlines suppressed the GSC differentiation defects, whereas germlines treated with the TOR inhibitor rapamycin had reduced levels of polyglutamine-containing proteins. Thus, ribosome biogenesis and ribosome levels can control stem cell differentiation via selective translation of CAG repeat-containing transcripts.

## Introduction

Understanding how stem cells self-renew and differentiate is crucial to understanding the mechanisms of development and disease (Cinalli et al., 2008; Morrison et al., 1997; Tang, 2012). Defects in ribosome biogenesis can impair stem cell differentiation and lead to diseases collectively called ribosomopathies (Armistead and Triggs-Raine, 2014; Barlow et al., 2010; Brooks et al., 2014; Calo et al., 2018; Higa-Nakamine et al., 2012; Mills and Green, 2017). Protein synthesis often increases during stem cell differentiation (Sanchez et al., 2016; Teixeira and Lehmann, 2019; Zhang et al., 2014), and inhibiting translation by modulating Target of Rapamycin (TOR) activity blocks terminal differentiation of various stem cells (Martin et al., 2022; Neumüller et al., 2008; Sanchez et al., 2016; Sun et al., 2010; Zhang et al., 2014). Nevertheless, how ribosome levels and translation control differentiation remains’ incompletely understood. In the ribosomopathy Diamond-Blackfan anemia (DBA), mutations in ribosomal proteins limit the pool of available ribosomes, which alters the translation of a select subset of transcripts in hematopoietic stem and progenitor cells, leading to impaired erythroid lineage commitment (Khajuria et al., 2018; Xue and Barna, 2012).

RNAs are extensively modified by post-transcriptional modifications (PTMs), including pseudouridylation (Granneman, 2004; Sloan et al., 2017; Tafforeau et al., 2013; Watkins and Bohnsack, 2012). The ribosomal RNA (rRNA) pseudouridine synthase subunit DKC1 is mutated in the ribosomopathy X-linked dyskeratosis congenita (X-DC), an inherited bone marrow failure syndrome that is sometimes associated with impaired neurodevelopment (Knight et al., 1999). DKC1 is a member of the snoRNA-guided H/ACA box, which deposits pseudouridine on rRNA at functionally important sites of the ribosome (Charette and Gray, 2000; Czekay and Kothe, 2021; Penzo and Montanaro, 2018). Mutations in DKC1 (Nop60B in *Drosophila*) can impair ribosomal binding to tRNAs and to internal ribosomal entry sites (IRES) from yeast to humans (Jack et al., 2011). Nevertheless, how H/ACA box dysfunction generates tissue-specific defects remains unclear.

During *Drosophila* oogenesis, differentiation of germline stems cells (GSCs) to an oocyte is sensitive to both ribosome biogenesis and translation (Blatt et al., 2020; Cinalli et al., 2008; Martin et al., 2022; Sanchez et al., 2016; Zhang et al., 2014). Oogenesis occurs in ovarioles beginning with the germline stem cells (GSCs) in the germaria (**Figure 1A**) (Lehmann, 2012; Xie and Spradling, 2000). The GSCs undergo asymmetric cell division to self-renew and give rise to daughter cells called cystoblasts (CBs) (Chen and McKearin, 2003b; Ohlstein and McKearin, 1997). The CB differentiates undergoing four incomplete mitotic divisions giving rise successively to 2-, 4-, 8-, and finally 16-cell cysts (Spradling et al., 1997; Xie and Spradling, 1998, 1998). One cell in the 16-cell cyst stage becomes the oocyte while the remaining 15 cells become the nurse cells that support the growing oocyte (**Figure 1A**) (Huynh and St Johnston, 2004; Kugler and Lasko, 2009; Lantz et al., 1994; Navarro et al., 2004). The GSCs and the CBs are marked by a round cytoskeletal structure called the spectrosome while the cysts are marked by a branched structure called the fusome (Chen and McKearin, 2003a; Ting, 2013). The 16-cell cyst becomes encapsulated by somatic cells to create an egg chamber that then goes through progressive development producing a mature egg (**Figure 1A**) (Huynh and St Johnston, 2004; Koch et al., 1967; Navarro et al., 2004).

**Figure 1:**
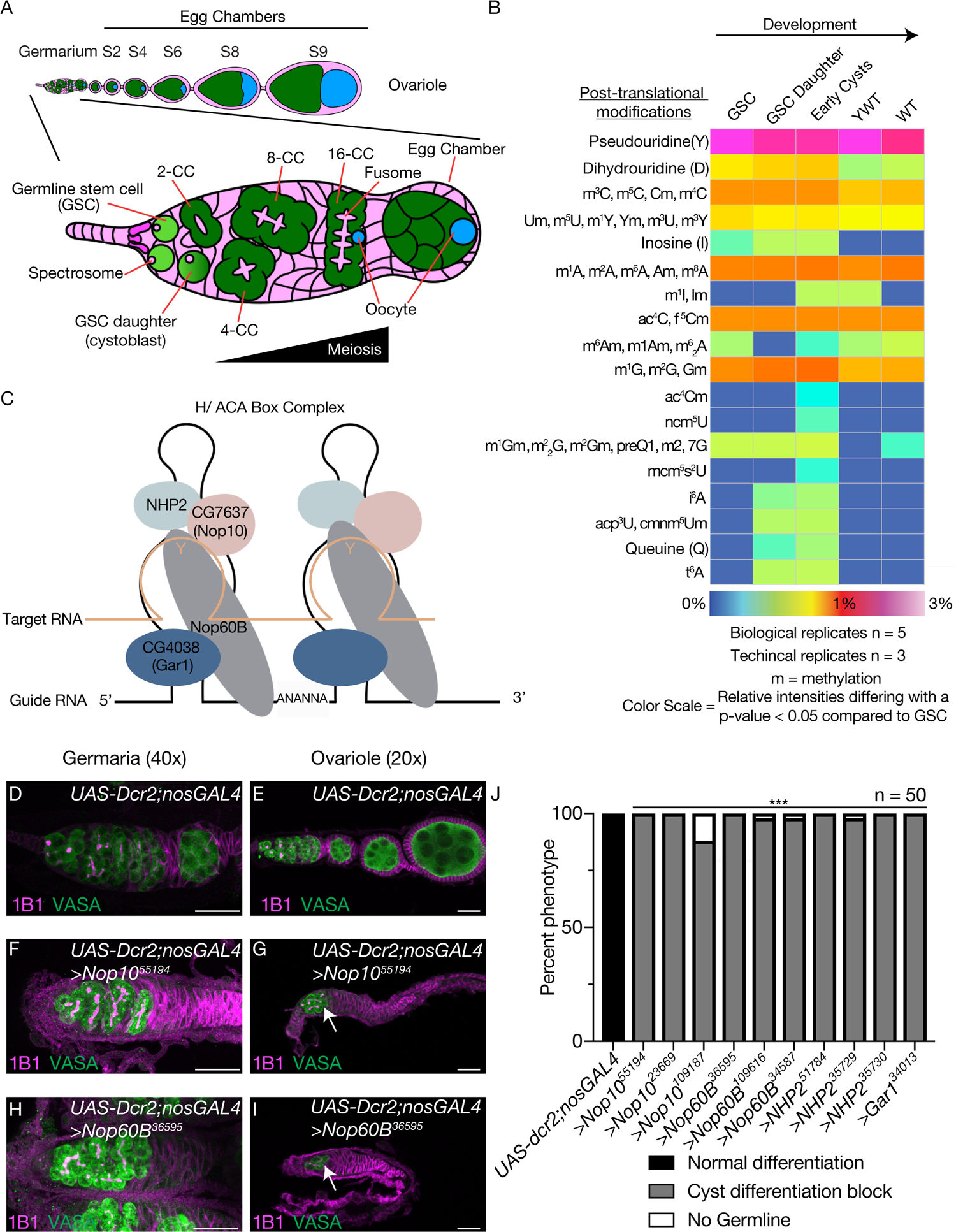
Pseudouridine is a critical modification required for oogenesis. (A) Schematic of a Drosophila ovariole and germarium. The germarium is present at the anterior tip of the ovariole and goes through successive stages of egg chamber development. The germline stem cells (GSC) (green) reside at the anterior tip of the germarium and are surrounded by somatic cells (magenta). The GSC divides to give rise to a GSC daughter or cystoblast (CB). The CB on the differentiation factor and undergoes incomplete mitotic divisions to give rise to a 2-,4-,8-, and 16-cell cyst (differentiating cysts). During the cyst stages, the germline transition from a mitotic fate to meiotic fate. The single cells are marked by spectrosomes (magenta) and the cysts are marked by the branched structure called fusomes (magenta). The 16-cell cyst buds off from the germarium and is encapsulated by the soma to generate an egg chamber. One of the 16-cells is designated as the oocyte (blue), going through successive egg chamber developmental eventually forming a mature egg. (B) Heat map of mass spectrometry analysis of RNA modifications obtained from total RNA extracts from germaria enriched for GSCs, GSC daughters, cysts, YWT, and Wild Type (WT). A heat map covers relative abundances up to 3%. The different colors express variations of relative abundance compared to the GSC with a p< 0.05 statistical significance. For each developmental stage 5 biological replicates were analyzed with 3 technical replicates of each biological replicates. (C) The H/ACA box is composed of four proteins CG4038 or Gar1 (dark blue), Nop60B (gray), NHP2 (light blue) and CG7637 or Nop10 (salmon). The H/ACA box uses a small RNA guide that corresponds to the target RNA where is complex deposits pseudouridine. (D, E) Images of 40x *UAS-Dcr2;nosGAL4* (driver control) germarium (D) and 20x *UAS-dcr2;nosGAL4* (driver control) ovarioles (E) stained with anti-1B1 (magenta) and anti-Vasa (green). (F, G) Images at 40x (F) and 20x (G) of germarium where *Nop10* is depleted in the germline and stained with anti-1B1 (magenta) and anti-Vasa (green) resulting in a cyst progression defect. White arrow marks cyst defect in germline depleted of H/ACA box members. Scale bar for all images is 20 µm. (H, I) Images at 40x (H) and 20x (I) of germarium where *Nop60B* is depleted in the germline and stained with anti-1B1 (magenta) and anti-Vasa (green) resulting in a cyst progression defect. White arrow marks cyst defect in germline depleted of H/ACA box members. Scale bar for all images is 20 µm. (J) Quantification of oogenesis defect phenotypes per genotype of H/ACA box germline depletion. Statistical analysis performed with Fisher’s exact test (n = 50 for all, *** p<0.0001).

In the CB, expression of Bag of marbles (Bam) promotes the progression from CB to an 8-cell cyst stage where expression of RNA-binding Fox protein 1 (Rbfox1) and Bruno 1 (Bru1) are required to specify an oocyte (Carreira-Rosario et al., 2016; Sugimura and Lilly, 2006). In parallel, several cells in the cysts initiate recombination mediated by the synaptonemal complex, which includes proteins such as crossover suppressor on 3 of Gowen (c(3)G), but only the specified oocyte commits to meiosis (Collins et al., 2014; Hughes et al., 2018; Page and Hawley, 2001). Rbfox1 is not only critical for female fertility but also for neurological functions (Carreira-Rosario et al., 2016; Gehman et al., 2012, 2011; Kucherenko and Shcherbata, 2018). Why some transcripts encoding differentiation factors, such as Rbfox1, are sensitive to ribosome levels is not known.

## Results

### RNA modifications are dynamic and essential for oogenesis

We aimed to identify dynamic RNA PTMs during oogenesis. Therefore, we enriched for five stages of oogenesis (1. GSCs; 2. GSC Daughter/CBs; 3. early-cysts; 4. germaria and early-stage egg chambers (young wild type (YWT)); 5. and late-stage egg chambers (WT)) which are critical milestones of germline development (Flora et al., 2018; McKearin and Spradling, 1990; Ohlstein and McKearin, 1997; Xie and Spradling, 1998). We performed tandem mass spectrometry on total RNA extracted from each of the enriched stages (**S1A-S1A’’’’**) (Flora et al., 2018; McKearin and Spradling, 1990; Ohlstein and McKearin, 1997; Xie and Spradling, 1998). For each enriched developmental stage, we performed 5 biological replicates, each with 3 technical replicates. We identified 18 groups of RNA PTMs represented by distinct mass:charge ratios, composed of 42 distinct RNA PTMs from a total of 172 known PTMs (**Figure 1B, Supplementary Table 1**). Pseudouridine, which is the most frequent PTM in RNA (Durairaj and Limbach, 2008), was the most abundant modification at all stages, followed by the monomethylations of the canonical RNA bases (**Figure 1B, Supplementary Table 1)**. Furthermore, we discovered a cohort of RNA PTMs, including inosine and dihydrouridine, that were not previously described during oogenesis (**Figure 1B, Supplementary Table 1**). Most RNA PTMs, including pseudouridine, were dynamic during GSC differentiation into an oocyte (**Figure 1B, Supplementary Table 1**).

To determine if the RNA modifications play a role in germline development, we performed an RNA interference (RNAi) screen utilizing a germline-specific *nanos-GAL4* driver to deplete RNA modifying enzymes in the germline, followed by immunostaining for Vasa, a germline marker, and 1B1, a marker of the cell membranes, spectrosomes and fusomes (Lasko and Ashburner, 1988; Zaccai and Lipshitz, 1996). We screened 33 unique genes annotated and predicted to be involved in RNA modification, and based on availability additional independent RNAi lines, for a total of 48 lines. Of the 33 distinct gene knockdowns, 2 resulted in loss of the germline, 14 in germaria defects, and 3 in egg chamber defects (**Supplementary Table 2**).

### The pseudouridine-depositing H/ACA box is required for oocyte specification

Among the genes whose knockown that caused defects in germaria, we found all four encoding components of the rRNA pseudouridine synthase, the H/ACA box: the catalytic subunit Nucleolar protein at 60B (Nop60B) and complex members CG7637 (Nop10), CG4038 (Gar1) and NHP2 (**Figure 1C, Supplementary Table 2**) (Giordano et al., 1999; Ni et al., 1997). Germline depletion of Nop60B in the background of an endogenous, GFP-tagged Nop60B reporter led to significantly reduced GFP levels in the germline (**Figure S1B-S1D**) (Sarov et al., 2016), verifying knockdown of *nop60B*. In addition, RT-qPCR analysis revealed significantly reduced levels of *Nop10* and *Nop60B* mRNAs upon germline knockdown (**Figure S1E**). Depletion of the H/ACA box components did not result in a germline viability defect, but rather to specific loss of GSCs and a cyst differentiation defect. Specifically, transition from 8-cell cyst stage to an egg chamber was blocked, as measured by the accumulation of 8-cell cysts (**Figure 1D-1J, S1F-S1P, Supplementary Table 2**) (Morita et al., 2018; Sanchez et al., 2016), which led to an absence of egg chambers and, in turn, sterility (**Figure S1Q**). Thus, the H/ACA box is required in the female germline for proper cyst differentiation.

We further investigated the role of the H/ACA box in cyst differentiation by analyzing control and H/ACA box germline-depleted ovaries carrying the differentiation reporter, BamGFP. We also stained ovaries for Vasa, 1B1, and the cyst-differentiation factors, Rbfox1 or Bru1 (Carreira-Rosario et al., 2016; Chen and McKearin, 2003b; Sugimura and Lilly, 2006). We found that cysts lacking H/ACA box members express BamGFP but have significantly reduced levels of Rbfox1 and Bru1 (**Figure 2A-G, S2A-S2D**). Moreover, cysts lacking H/ACA box components did not specify an oocyte, as cysts were devoid of localized Egalitarian (Egl), the oocyte determinant, and exhibited reduced expression of the synaptonemal complex component C(3)G (**Figure S2E-S2L**) (Anderson et al., 2005; Carpenter, 1994; Huynh and St Johnston, 2000; Mach and Lehmann, 1997; Page and Hawley, 2001), consistent with a differentiation block.

**Figure 2:**
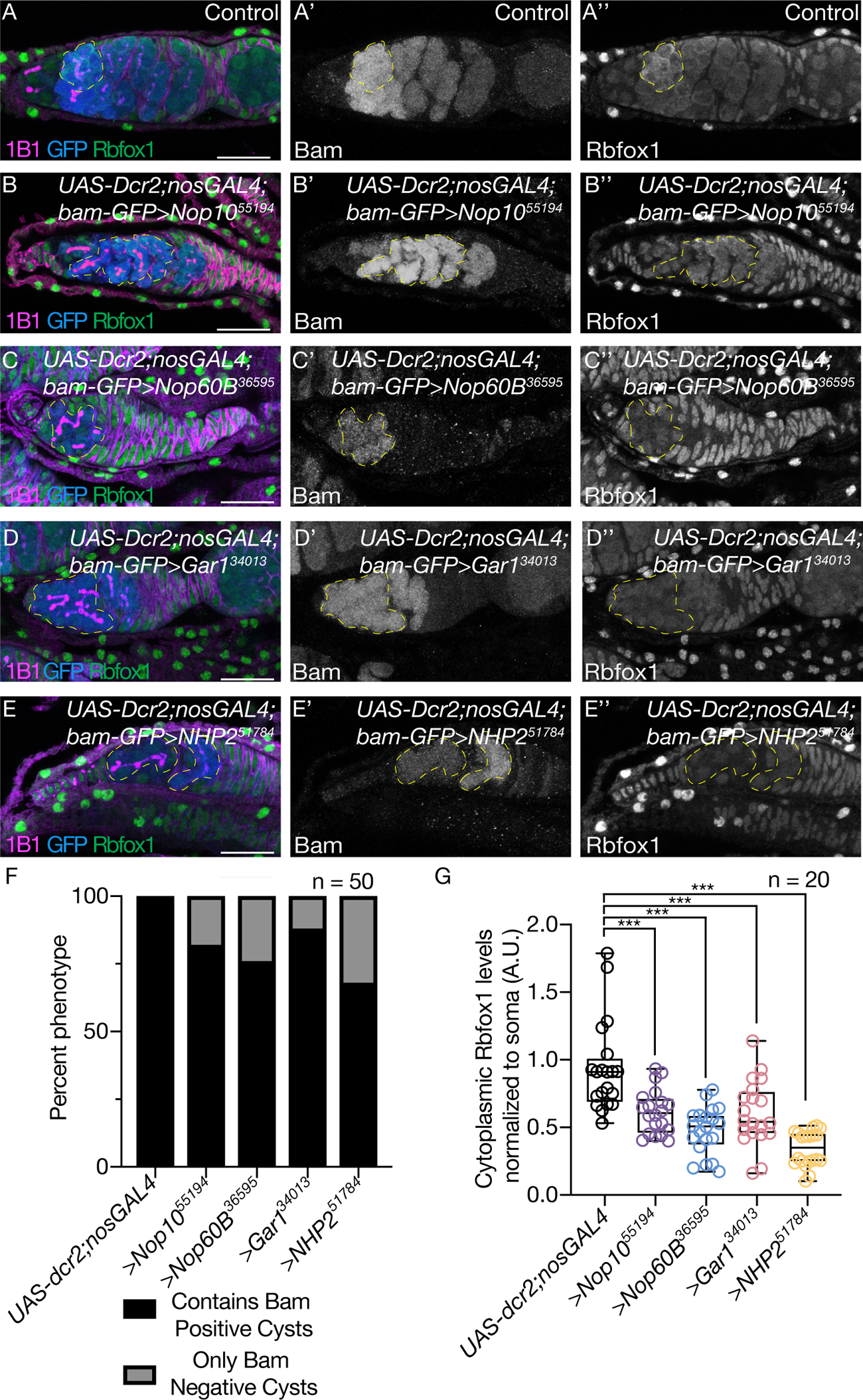
The H/ACA box is required for proper cyst differentiation and meiotic progression. (A-E”) *UAS-Dcr2;nosGAL4;bam-GFP* (driver control) germaria (A) and germline depletion of *Nop10* (B), *Nop60B* (C), *Gar1* (D), and *NHP2* (E) stained with anti-1B1 (magenta), anti-GFP (blue) and anti-Rbfox1 (green). GFP (A’, B’, C’, D’, E’) and Rbfox1 (A’’, B’’, C’’, D’’, E’’) are shown in gray scale. Yellow dotted lines outline cysts that are positive for GFP but have lower Rbfox1 levels for all images. Scale bar for all images is 20 µm. (F) Quantification of oogenesis defect phenotypes per genotype. Loss of the H/ACA box results in Bam positive cysts. Statistical analysis performed with Fisher’s exact test (n = 50 each, *** p<0.0001). (G) Quantification of cytoplasmic Rbfox1 levels normalized to soma in germline depleted of *Nop10*, *Nop60B*, *Gar1* or *NHP2*. Loss of H/ACA box results in lower levels of Rbfox1 levels. Statistics performed were Dunnett’s multiple comparisons test post-hoc test after one-way ANOVA (n = 20 each, *** p<0.0001).

To determine the specific stage of oogenesis that requires H/ACA box activity, we first characterized the expression of the endogenous, GFP-tagged Nop60B reporter (Sarov et al., 2016). Nop60B-GFP levels increased from the cyst stages to early egg chambers (**Figure S3A-S3B**). Utilizing a pseudouridine antibody, we observed a corresponding increase in pseudouridine levels from the 8-cell cyst to the newest egg chamber and this increase depended on the H/ACA box (**Figure S3C-S3G**). Given these observations, and that loss of H/ACA box components resulted in an accumulation of 8-cell cysts (**Figure S1P**) (Morita et al., 2018), we hypothesized that the H/ACA box is required in the cysts for the transition into an oocyte. To test this, we depleted *Nop60B* and *Nop10* in the cysts utilizing a *bamGAL4* driver, which is active in the 2-8 cell cyst stages. We observed an accumulation of cysts with significantly reduced levels of Rbfox1 (**Figure S4A-S4H**) (Carreira-Rosario et al., 2016; Chen and McKearin, 2003b). Taken together, these data suggest that the H/ACA box is required in the cyst stages for differentiation into an oocyte.

### The H/ACA box promotes ribosome biogenesis and the translation of differentiation factors during oogenesis

The primary activity of the H/ACA box is to deposit pseudouridine on rRNA, thereby promoting ribosome biogenesis in the nucleolus (Gilbert, 2011; Kiss et al., 2010; Ni et al., 1997; Omer et al., 2000). Nop60B-GFP colocalized with Fibrillarin in the nucleolus as previously observed (Ochs et al., 1985) (**Figure S5A-S5B**). In addition, loss of Nop10 and Nop60B resulted in cysts with hypertrophic nucleoli compared to wild-type cysts, suggesting a ribosome biogenesis defect (**Figure S5C-S5F**) (Neumüller et al., 2008; Sanchez et al., 2016). To verify that the H/ACA box deposits pseudouridine on rRNA during oogenesis, we co-immunopurified the 40S and 60S ribosomal subunits from the germline, utilizing a germline-enriched HA-tagged ribosomal protein RpS5b (**Figure S5G-S5I**) (Jang et al., 2021) (Chen and Dickman, 2017). Mass spectrometry analysis showed that loss of the H/ACA box member Nop10 led to a significant decrease of pseudouridine on rRNA relative to controls (**Figure 3A, Supplemental Table 3**). In addition, we observed a decrease in both the 40S and 60S subunits and in polysomes of Nop60B-depleted ovaries as compared to controls (**Figure S5J**) (Cheng et al., 2019), suggesting a ribosome biogenesis defect upon loss of the H/ACA box. Thus, consistent with previous findings, the H/ACA box deposits pseudouridine on rRNA to promote ribosome biogenesis in the germline.

**Figure 3:**
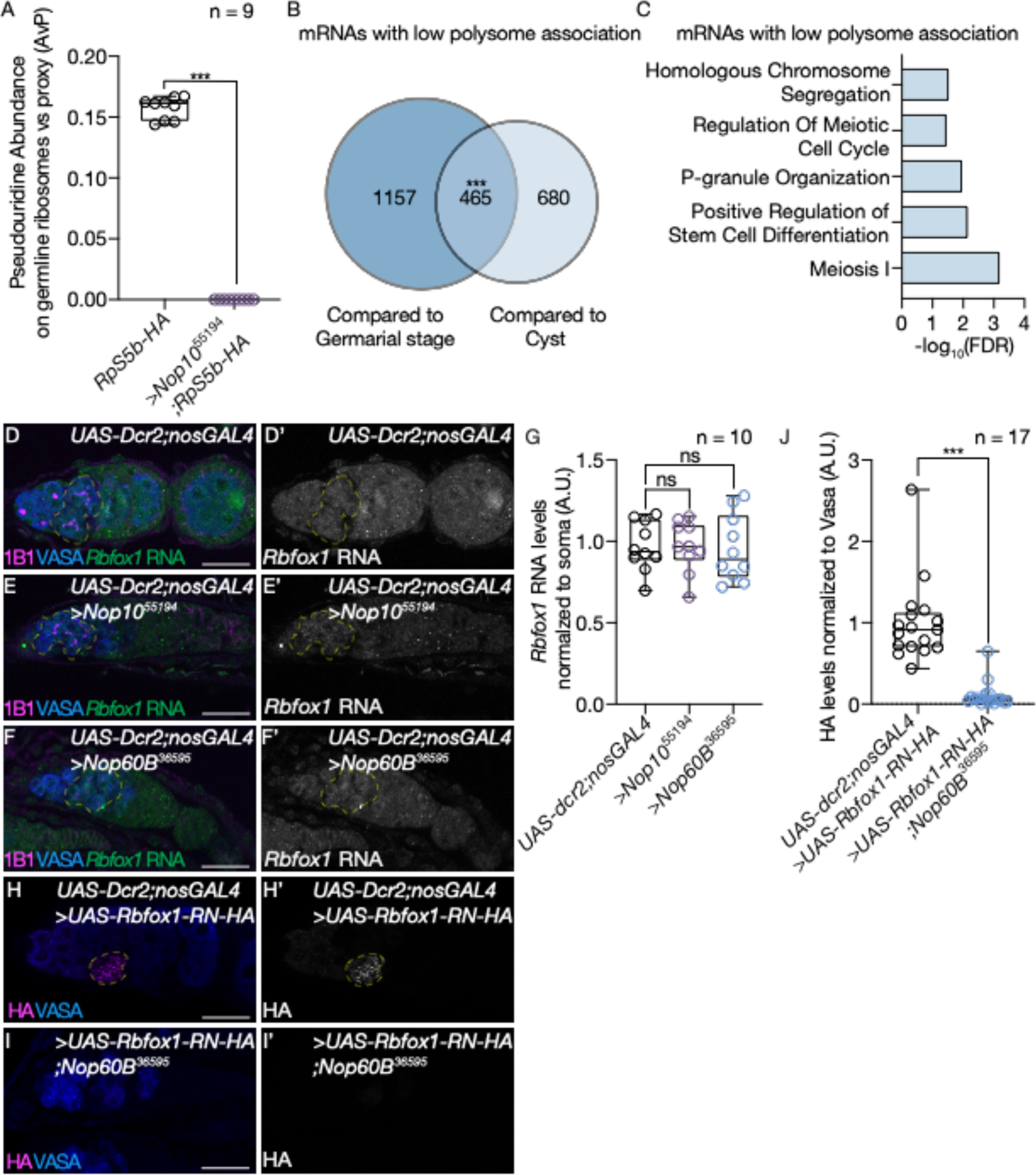
The H/ACA box is required for translation of meiotic mRNAs. (A) Mass spectrometry analysis of rRNA isolated from germline ribosomal pulldowns showing reduced pseudouridine levels on rRNA. Statistics performed were t-test of pseudouridine levels comparing germaria enriched for cysts and *Nop10* depleted germaria. For each developmental stage at least 2 biological replicates were analyzed with 3 technical replicates of each biological replicates (*** p<0.0001). (B) Venn diagram illustrating overlap of Nop60B-polysome <-2 fold less association with the ribosome upon loss of *Nop60B* (n = 2, e < 2.87 × 10^-192^, Hypergeometric Test). Controls were cysts, enriched through heat shock, and young wild-type (YWT), which includes germarial stages and a few egg chambers. (C) Significant biological process GO terms of shared lowly associated mRNAs in ovaries depleted of *Nop60B* compared to control sets, showing an enrichment for mRNAs associated with meiosis 1. (D) In situ hybridization to *Rbfox1* RNA (green) and staining with anti-1B1 (magenta) and anti-Vasa (blue) in *UAS-Dcr2;nosGAL4* (driver control) germaria and (E,F) germline-depleted of *Nop10* and *Nop60B*. *Rbfox1* RNA is shown in gray scale (D’, E’, F’). Yellow dotted line outlines *Rbfox1* RNA. (G) Quantification of *Rbfox1* RNA levels in germline depleted of varying members of *Nop10* and *Nop60B* normalized to soma showing no significant difference in Rbfox1 levels. Statistics performed were Dunnett’s multiple comparisons test post-hoc test after one-way ANOVA (n = 10 for all, not significant, P>0.9999 and p=0.9792 respectively). (H) Germaria of *UAS-Dcr2;nosGAL4* (driver control) driving UAS-Rbfox1-RN-HA and (I) germline depleted of *Nop60B* driving Rbfox1-RN-HA. Germaria stained with anti-Vasa (blue) and anti-HA (magenta). HA is shown in gray scale (H’, I’). (J) Quantification of HA levels in control vs germline depleted of *Nop60B* normalized to vasa showing reduced Rbfox1-RN-HA levels in the germline. Statistics performed were unpaired t-test (n = 17 for all, *** p<0.0001). Yellow dotted line outlines Rbfox1-RN-HA.

To test if loss of the H/ACA box and consequent aberrant ribosome biogenesis affects mRNA translation during oogenesis, we performed polysome-seq of ovaries depleted of Nop60B in the germline and of gonads enriched for cysts stages (Flora et al., 2018; Martin et al., 2022; Ohlstein and McKearin, 1997). As enriching for early cyst stages includes a heat shock step (see methods), we also analyzed early-stage egg chambers to control for heat shock effects. We detected 465 mRNAs with a reduced polysome association in *Nop60B*-knockdown versus the controls, whereas 638 mRNAs showed an increased polysome association (**Figure 3B, S6A-S6C**). These data suggest that the H/ACA box regulates the synthesis of a cohort of proteins. GO term analysis revealed that mRNAs with an elevated polysome association were enriched in the mitotic cell cycle, whereas those with reduced polysome association included factors that promote meiosis 1, meiotic cell cycle and homologous chromosome segregation (**Figure 3C, S6D**), such as the synaptonemal complex members C(3)G and Corona (cona), consistent with reduced C(3)G protein levels upon depletion of the H/ACA box (**Figure S2I-S2L**).

The levels of *Rbfox1* and *Bru1* mRNAs were not significantly reduced in the germline upon depletion of the H/ACA box, as indicated by fluorescent *in situ* hybridization (**Figure 3D-3G, Figure S6E-S6H**). To determine if the H/ACA box is required for translation of Rbfox1 and Bru1, we overexpressed Rbfox1 or Bru1 under the control of UAS/GAL4 system. Rbfox1 and Bru1 proteins were detected in the control germaria, but not in the H/ACA box-depleted germaria (**Figure 3H-3J, S6I-S6K**) (Carreira-Rosario et al., 2016; Filardo and Ephrussi, 2003), suggesting that their translation is impaired upon loss of the H/ACA box.

We considered that the H/ACA box is required for oogenesis due to its role in ribosome biogenesis. This hypothesis predicts that compromised ribosome biogenesis will phenocopy loss of H/ACA box components. We impaired ribosome biogenesis by depleting ribosomal protein paralogs RpS10b and RpS19b in the germline, as the depletion of other ribosomal proteins that do not have paralogs results in GSC differentiation defects or loss of cyst stages that would mask the cyst differentiation block (Jang et al., 2021; Martin et al., 2021; McCarthy et al., 2022; Sanchez et al., 2016). Depletion of RpS10b and RpS19b phenocopied loss of H/ACA box components, leading to a block in cyst differentiation and decreased levels of Rbfox1 and Bru1 proteins without a concomitant loss of their mRNAs (**Figure S7A-S7K, S8A-S8H**). The H/ACA box can also pseudouridylate mRNAs and tRNAs (Czekay and Kothe, 2021). However, immunopurification of pseudouridine did not enrich for the mRNAs with perturbed translation upon loss of the H/ACA box (**Supplemental Table 4**), suggesting that these targets are not directly pseudouridylated. In addition, whereas the loss of the H/ACA box blocks cyst differentiation, loss of tRNA pseudouridylation enzymes results in a different phenotype – loss of cyst stages (**Figure S9A-S9J**). Thus, our data suggest that the H/ACA box and pseudoruridinylation are important for ribosome biogenesis during oogenesis.

### H/ACA box-dependent differentiation factors are PolyQ proteins

To identify shared properties among the mRNAs with reduced polysome association upon loss of the H/ACA box, we performed a motif analysis of the 5’UTRs, CDS and 3’UTRs of this subset of mRNAs compared to a set of control mRNAs. We observed a motif of repeating CAG nucleotides that was highly enriched in the CDS of downregulated transcripts compared to the control unregulated mRNAs (**Figure 4A**). In addition, we found motifs that were enriched in the in the 3’UTRs or 5’UTRs of downregulated transcripts, albeit in a smaller subset of RNAs (**Supplementary Table 5**). The CAG motifs in the downregulated transcripts were in frame, such that the encoded proteins are highly enriched in glutamine (Q) over a region of 21 amino acids (**Figure 4B**). Indeed, Rbfox1 and Bru1 both contain such repeating CAG motifs in the mRNA and polyQ in the protein (**Figure S10A-S10D**) (Kucherenko and Shcherbata, 2018).

**Figure 4:**
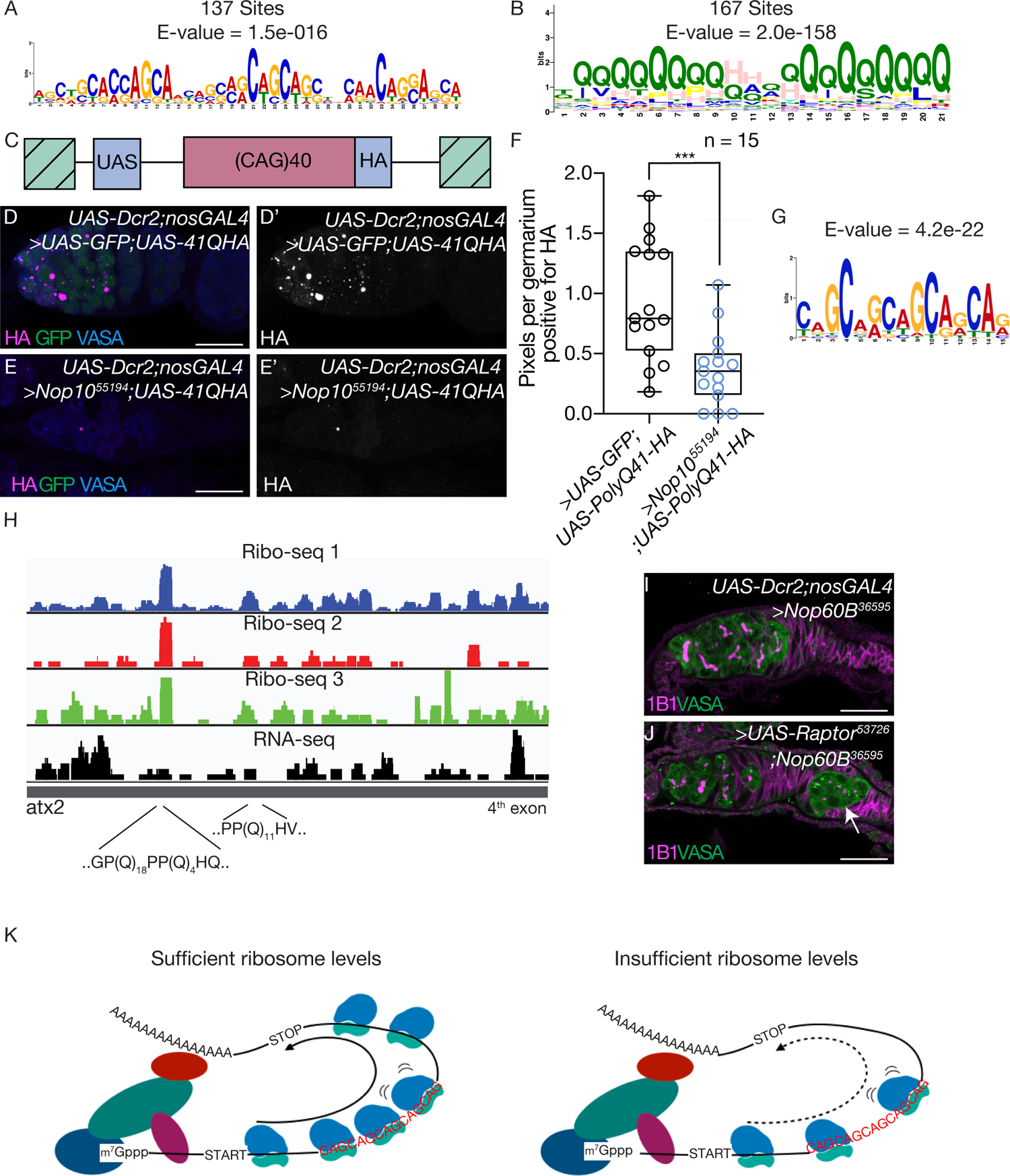
The H/ACA box is required for translating polyQ proteins. (A) Repeating CAG motif identified by MEME enriched in the CDS of 137 mRNAs that are lowly associated with polysomes in germaria depleted of *Nop60B*. (B) Repeating polyQ motif present in 167 sites identified by MEME enriched that are present in the mRNAs that are lowly associated with the ribosome in germaria depleted of Nop60B. (C) Schematic of CAG reporter which codes for a 41Q protein with an HA tag and which can be driven by the *UAS-GAL4* in a tissue specific manner. (D) Confocal image of poly41Q-HA reporter driven in *UAS-Dcr2;nosGAL4* controls flies or poly41Q-HA reporter driven in *Nop10* depleted germaria (E) stained with anti-HA (magenta), anti-GFP (green) and anti-Vasa (blue). UAS-GFP was also driven to ensure equal GAL4 dosage. HA is shown in gray scale (D’ and E’). Scale bar for all images is 20 µm. (F) Quantification of percent of pixels per area of HA in control vs germline depleted of *Nop10* showing a reduction of HA signal in *Nop10* depleted germaria. Statistics performed were unpaired t-test (n = 15 for all, *** p=0.0007). (G) Repeating CAG motif identified by MEME at peak regions in mRNAs detected by ribosome footprinting. (H) Ribosome footprint distribution on *atx2* mRNA, illustrating a peak in exon 4 found in common among the 3 Ribo-Seq libraries (blue, red and green) but not in the RNA-Seq library (black). A polyQ stretch is present at the ribosome peak. (I, J) Germaria depleted for Nop60B (I) or Nop60B (J) while simultaneously overexpressing Raptor-HA using *UAS-Dcr2;nosGAL4*. Germaria stained with anti-Vasa (green) and anti-1B1 (magenta). Arrow points at egg chamber. (>Nop60B RNAi, N = 91, 1.1% contained 1^st^ egg chamber while >Nop60B RNAi;UAS-Raptor, N = 151, 9.9% contained 1^st^ egg chamber, Fisher’s exact test, p=0.0037). (K) Representative model showing that a sufficient level of ribosomes is required for translation of meiotic mRNAs containing a repeating CAG motif. Proper ribosome levels allow for translation of these mRNAs to promote terminal differentiation. Ribosome insufficiency result in reduced translation of meiotic mRNAs, due to ribosome stalling or slowing, that contain the repeating CAG motif causing a failure of terminally differentiate.

To determine if the H/ACA box is required to translate CAG repeat-containing mRNAs during oogenesis, we expressed a CAG reporter encoding an HA-tagged, polyQ protein, that was previously used to model polyglutamine toxicity from Huntington’s disease (**Figure 4C**) (Fayazi et al., 2006). We co-expressed the CAG reporter with GFP in the control, to ensure equal GAL4 dosage, and the CAG reporter in Nop10-depleted germlines. We found that loss of the H/ACA box specifically reduced the levels of the polyQ protein (**Figure 4D-4F, S10E-S10F).** Furthermore, depletion of RpS19b and Nop60B also resulted in a significant decrease in polyQ protein accumulation, but loss of Nop60B did not significantly alter the levels of GFP **(Figure S10G-S10M).** Thus, the H/ACA box and ribosome biogenesis are required for translation of polyQ-containing proteins.

### CAG repeat regions show increased density of ribosomes

One proposed mechanism of polyQ expansion-induced defects is the disruption of translation by ribosome stalling (Eshraghi et al., 2021). To determine if the H/ACA box affects elongating ribosomes on, and hence translation of, mRNAs encoding polyQ proteins, we performed ribosome footprinting (Ribo-Seq). Because we were unable to acquire sufficient material from H/ACA box-depleted germaria, we utilized ovaries enriched for late-stage oocytes (late-ovaries), which have reduced pseudouridine levels but can be obtained in sufficient quantity (**Figure 1B, Supplementary Table 1)**. Three late-ovary Ribo-Seq libraries were generated, each with a corresponding RNA-Seq library. Correlation analysis showed consistent and reproducible ribosome footprint distributions among the three Ribo-Seq libraries (Pearson r > 0.9 for all comparisons; **Supplemental Table 6)**. We hypothesized that stalled ribosomes might result in local enrichment of ribosome footprints. We therefore sought to identify peaks in the Ribo-Seq data across the transcriptome. Our peak detection and subsequent motif analysis identified 123 mRNAs containing at least one CAG-rich segment within 30 nucleotides of a ribosome footprint peak in at least 2 of the 3 Ribo-Seq libraries that were not present in the RNA-seq libraries (**Figure 4G, Supplemental Table 6**). 46 of the 123 identified mRNAs encode at least one polyQ tract (≥ 4 consecutive CAG codons) near ribosome footprint peaks (**Figure 4H, Supplemental Table 6**). This motif is highly reminiscent of the motif identified in the mRNAs with low polysome association in H/ACA box depleted germlines **(Figure 4A)**.

The motif identified by Ribo-Seq contains 5 CAGs in a row **(Figure 4A** and **4G)**. To determine if the 5-CAG motif is overrepresented in the set of target mRNAs with low polysome association, we performed a Find individual Motif Occurrence (FIMO) analysis. We found that 181 out of 465 (39%) targets contain a significant motif representative of the 5-CAG motif including *bru1* and *C(3)G* **(Figure S2A-S2D, S2I-S2K, Supplemental Table 7)**. We infer that CAG repeat sequences have high ribosome density and are present in mRNAs whose translation is sensitive to reduced ribosome biogenesis during oogenesis.

### Tor signaling partially restores differentiation and modulates polyQ translation

The Target of Rapamycin (TOR) pathway is a critical positive regulator of ribosome biogenesis (Wullschleger et al., 2006; Yerlikaya et al., 2016). To further determine if loss of the H/ACA box blocks cyst differentiation due to reduced ribosome biogenesis, we increased ribosome biogenesis by overexpressing the TORC1 co-factor, Raptor, in the H/ACA box-depleted germline (Martin et al., 2006; Wang et al., 2012). We observed a partial yet significant alleviation of the cyst differentiation defect, such that an egg chamber was formed **(Figure 4I-4J)**. We next asked if decreasing TOR activity and, in turn, ribosome biogenesis could diminish expression of the polyQ reporter. Specifically, flies expressing the germline CAG reporter were treated with the inhibitor of mTOR, Rapamycin, and displayed lower levels of germline polyQ when compared to controls **(Figure S10N-S10P**). Thus, our data suggest that modulating ribosome levels via the Tor pathway can effectively regulate translation of polyQ-containing proteins.

## Discussion

We used the power of *Drosophila* genetics combination with mass spectrometry to determine the developmental profile of RNA PTMs and identify a cohort of PTMs that are required for proper oogenesis. Specifically, we found that pseudouridine abundance is dynamic and regulated by the H/ACA box, a pseudouridine synthase, and is required for proper cyst differentiation and oocyte specification. Using polysome-seq analysis, we found that CAG repeat mRNAs encoding polyQ-containing proteins have reduced polysome association upon loss of the H/ACA box. These polyQ proteins include germ cell differentiation and meiosis promoting factors such as Rbfox1 and Bru1. Moreover, we found that CAG repeat regions accumulate ribosomes, potentially acting as a ribosome sink. Taken together, our data suggest under condition of low ribosome levels, the CAG repeat containing regions can impede proper translation by sequestering ribosomes internally causing translation of polyQ-containing proteins to be sensitive to ribosome levels **(Figure 4K)**. Taken together, we find that the H/ACA box promotes ribosome biogenesis during oogenesis and, in turn, the translation of CAG repeat mRNAs required for differentiation **(Figure 4K)**.

Ribosomapathies predispose individuals to neurological deficits, but the etiology of this defect is unclear (Aspesi and Ellis, 2019; Cheng et al., 2019; Mills and Green, 2017). Neurons express and require several polyQ-containing proteins, including Rbfox1 (Gehman et al., 2012, 2011; Kucherenko and Shcherbata, 2018). We find that the translation and levels of Rbfox1 are sensitive to ribosome levels during oogenesis (McCarthy et al., 2022). By extension, neuronal deficits observed in ribosomapathies could be due to inability to translate critical polyQ-containing proteins in neurons.

While polyQ stretches facilitate phase transition, large CAG expansion and polyQ protein aggregates are associated with diseases such as Huntington’s disease (Adegbuyiro et al., 2017; Sugars and Rubinsztein, 2003). A genome-wide association study revealed that the onset of Huntington’s disease is due to large expansions of CAG repeats and is accelerated by DNA repair genes as well as E3 ubiquitin protein ligase (UBR5) (Lee et al., 2019). In embryonic stem cells, UBR5 has been shown to physically interact with the H/ACA box to promote rRNA maturation suggesting that these factors could collaborate to contribute to early onset of Huntington’s Disease (Saez et al., 2020). Furthermore, Huntington’s Disease mouse models have shown CAG expansions induce ribosome stalling by impeding ribosome translocation, thereby inhibiting protein synthesis (Eshraghi et al., 2021). These data together with our findings that, during development, the H/ACA box promotes translation of CAG repeat containing RNAs suggest that translation dyregulation could be a key feature of CAG expansion diseases. While the early onset is determined by CAG length, translation of CAG repeats into polyQ proteins can cause protein aggregation and toxicity (Bates, 2003; Lee et al., 2019; Ross and Poirier, 2004). Our finding that translation of such polyQ proteins is sensitive to ribosome levels reveal new potential therapeutic targets. For instance, several TOR inhibitors have been generated to primarily treat cancers; the mechanism we have identified provides a potential pathway to repurpose these drugs to reduce polyQ protein aggregation in various repeat expansion disease states (Noda, 2017; Ravikumar et al., 2004; Wyttenbach et al., 2008; Yee et al., 2021).

## Acknowledgements

We are grateful to all members of the Rangan laboratory, Dr. Marlow, Dr. Siekhaus and Life Science Editors for discussion and comments on the manuscript. We would like to thank Lehmann lab for Egl antibody, Dr. Teleman for p-S6 antibody, Bloomington Drosophila Stock Center, Vienna Drosophila Resource Center and FlyBase for fly stocks and reagents. Furthermore, we would like to thank CFG Facility at the University at Albany (UAlbany) for performing RNA-seq analyses. P.R. is funded by NIH/NIGMS (RO1GM11177 and RO1GM135628). E.R.G. is funded by NIH/NIGMS (R35 GM126967) and D.F. is funded by by NIH/NIGMS (R01 GM123050 and R01 GM121844) and NIH/NIDA (R01 DA04611).

## Materials and Methods

### Fly lines

Flies were grown at 25-29°C and dissected between 0-3 hrs and 1-3 days post-eclosion. Heat shock experiments were performed on 1 day old flies. The following RNAi stocks were used in this study: *nsun2 RNAi* (Bloomington #62495), *Trm7-34 RNAi* (Bloomington #62499), *CG32281 RNAi* (Bloomington #51764), *rswl* (Bloomington #44494), *CG9386* (Bloomington #33364), *Nsun5* (Bloomington #32400), *CG11109* (Bloomington #56897), *CG11447* (Bloomington #43207), *CG3021* (Bloomington #55144), *RluA-2* (VDRC #34152), *RluA-2* (VDRC #106382), *Wuho* (Bloomington #61281), *CG10903* (VDRC #57481), *RluA-1* (VDRC #41757), *RluA-1* (VDRC #41758), *RluA-1* (VDRC #109586), *Tailor* (Bloomington #36896), *Pus1* (Bloomington #53288), *NOP60B* (Bloomington #36595), *CG6745* (Bloomington #41825), *CG7637* (Bloomington #55194), *NHP2* (Bloomington #51784), *CG4038* (Bloomington #34013), *CG34140* (Bloomington #38951), *CG34140* (Bloomington #57311), *CG3645* (VDRC #107156), *CG1434* (VDRC #104876), *CG3434* (VDRC #45130), *tgt* (VDRC #41644), *AlkB* (Bloomington #43300), *Paics* (Bloomington #62241), *Ras* (Bloomington #31654), *Ras* (Bloomington #51717), *Ras* (Bloomington #31653), *pfas* (Bloomington #36686), *pfas* (Bloomington #80831), *bam RNAi; hs-bam (*Ohlstein, B. & McKearin, D), RpS10b (Bloomington #43976), RpS19b (VDRC #22073), UAS-raptor-HA (Bloomington #53726).

The following tissue-specific drivers were used in this study: *UAS-Dcr2;nosGAL4* (Bloomington #25751), *UAS-Dcr2;nosGAL4;bamGFP* (Lehmann Lab), *If/CyO;nosGAL4* (Lehmann Lab), *nosGAL4;MKRS/TM6* (Bloomington #4442), and *TjGAL4/CyO* (Lehmann Lab).

The following stocks were used in this study: *RpS5b-HA* and *UAS-Rbox1-RN* (Buszazak Lab) and *UAS-Tkv* (Bloomington#36536), *UAS-Rbox1-RN* (Buszazak Lab) *UAS-Bruno* (Ephrussi Lab) and UAS-41Q.HA (Bloomington #30540).

### Rapamycin treatment

One day prior to treatment, 400 μL of 100uM Rapamycin or 400μL of ethanol was added to the top of food and allowed to dry. Flies were crossed at 18°C and collected 1-2 days post-eclosion. Flies were placed on food and temperature shifted to 29°C. Every other day flies were placed onto fresh food with 400 μL of 100uM Rapamycin or 400μL of ethanol for a total of 7 days. Flies were dissected as described below.

### Genotypes used to enrich specific stages of germline

To enrich for germline Stem Cells: *nosGAL4>UAS-tkv (Flora et al., 2018; Xie and Spradling, 2000, 1998)*. Cystoblasts: *nosGAL4>bam RNAi* (Chen and McKearin, 2003a, 2003b; McKearin and Spradling, 1990). Differentiating Cysts: *nosGAL4>bam RNAi; hs-bam* (McCarthy et al., 2022; Ohlstein and McKearin, 1997). Female flies were heat shocked at 37° C for 2 hours, incubated at room temperature for 4 hours and heat shocked again for 2 hours. This was subsequently repeated the next day and flies were dissected. Young Wild Type: Female flies were collected and dissected within 2 hours of eclosion. To dissect wild-type ovaries, 2-3 day old females (*UAS-dcr;nosGAL4*) were fatten overnight and dissected the next day.

### Dissection and Immunostaining

Ovaries were dissected into 1X PBS and fixed for 10 minutes in 5% methanol-free formaldehyde (Flora et al., 2018). Samples were washed in 1 mL PBT (1X PBS, 0.5% Triton X-100, 0.3% BSA) 4 times for 10 minutes each. Primary antibodies were added in PBT and incubated at 4°C rotating overnight. Samples were washed 3 times for 10 minutes each in 1 mL PBT, and once in 1 mL PBT with 2% donkey serum (Sigma) for 10 minutes. Secondary antibodies were added in PBT with 4% donkey serum and incubated at 4°C rotating overnight. Samples were washed 4 times for 10 minutes each in 1 mL of 1X PBST (0.2% Tween 20 in 1x PBS). Vectashield with DAPI (Vector Laboratories) was added for 30 minutes before mounting. The following primary antibodies were used: mouse anti-1B1 (1:20, DSHB), rabbit anti-Vasa (1:1000, Rangan Lab), chicken anti-Vasa (1:1000), rabbit anti-GFP (1:2000, Abcam, ab6556), rabbit anti-Egl (1:1000, Lehmann Lab), mouse anti-pseudouridine (1:1000, MBL Life Sciences), mouse anti-C3G (1:1000, Hawley Lab), rat Anti-HA(1:500, Roche, 11 867 423 001), mouse anti-Fibrillarin (1:50, Fuchs Lab), guinea pig anti-Rbfox1 (1:1000, Buszazak Lab) and rabbit anti-phosphorylated-S6 (1:200, Teleman Lab).The following secondary antibodies were used: Alexa 488 (Molecular Probes), Cy3, and Cy5 (Jackson Labs) were used at a dilution of 1:500.

### Fluorescence Imaging

The tissues were visualized under 20X oil and 40X oil objective lenses and images were acquired using a Zeiss LSM-710 confocal microscope. Confocal images were processed with ImageJ. The images were quantified using ImageJ with the Measurement function.

### AU quantification of protein or in situ

To quantify antibody staining intensities for Rbfox1, Bruno, GFP, and pseudouridine or in situ probe fluorescence in germ cells, images for both control and experimental germaria were taken using the same confocal settings. Z stacks were obtained for all images. Similar planes in control and experimental germaria were chosen, the area of germ cells positive for the proteins or in situs of interest was outlined and analyzed using the ‘analyze’ tool in Fiji (ImageJ). The mean intensity and area of the specified region was obtained. An average of all the ratios (Mean/Area), for the proteins or in situs of interest, per image was calculated for both, control and experimental. Germline intensities were normalized to somatic intensities or if the protein or in situ of interest is germline enriched and not expressed in the soma they were normalized to Vasa or background. The highest mean intensity between control and experimental(s) was used to normalize to a value of 1 A.U. on the graph.

To quantify polyQ-HA, images were first filtered with a median pixel of 1. The program set the threshold values using max entropy threshold for the images and the outline of the germline was traced using the germline marker Vasa. The percent pixel count per the germline area was found and normalized to the highest mean intensity between control and experimental(s). For rapamycin treatment, 15 control and 15 treated germaria were used. Three randomly selected slices of each stack (total of 45 slices) were quantitated for both control and rapamycin treated germaria.

### Egg Laying Assay

Egg laying assays were conducted in triplicate in vials containing standard fly food. Prior to the assay, dry yeast was added to each vial along with 3 adult females (all 1 day post-eclosion) and 1 male. Flies were incubated at 29°C overnight. The flies were then placed in a new tube and the total number of eggs counted.

### RNA Isolation

Ovaries were dissected in 1X PBS and homogenized by motorized pestle in 100μL of TRIzol (Invitrogen, 15596026). RNA was isolated by adding an additional 950 μL of TRIzol and 200uL of chloroform with mixing. Samples were centrifuged at 13,000 rpm, 4°C for 15 minutes. The aqueous phase was transferred to a new tube, nucleic acids were precipitated using 1 mL of 100% ethanol, 52 μL of 3M sodium acetate and precipitated for >1 hour at −20°C. Samples were centrifuged at 13,000 rpm, 4°C for 20 minutes. Ethanol was decanted, pelletsw were washed twice with 1 mL of 70% ethanol and dried at room temperature for 10 minutes. Pellets were dissolved in 20 μL RNase free water and placed in a 42°C for 10 minutes. The concentration of nucleic acid samples were measured on a spectrophotometer. The samples were treated with DNase (TURBO DNA-free Kit, Life Technologies, AM1907) and incubated at 37°C for 30 min. DNAse was inactivated using the included DNAse. Inactivation reagent and buffer according to manufacturers instructions.

### RNA-seq and Polysome-seq library preparation

RNA was isolated as previously described above. Total RNA samples were run on a 1% agarose gel to assess sample integrity (McCarthy et al., 2018). To generate RNA-seq libraries, total RNA was incubated with poly(A) selection bead. mRNA libraries were prepared using the NEXTflex Rapid Directional RNAseq Kit (BioO Scientific Corp.). Fragmentation of the mRNA was achieved by incubating 95°C for 13 minutes to produce ∼300 bp fragments. Single-end mRNA sequencing (75 base pair) was performed for each sample with an Illumina NextSeq500, carried out by the Center for Functional Genomics (CFG). The sequenced reads were aligned to the *Drosophila melanogaster genome* (UCSCdm6) using HISTAT2 with Refseq annotate transcripts as a guide. featureCounts was used to generate raw counts and differential gene expression was assayed by Deseq, using a false discovery rate of (FDR) of 0.05, and genes with 2-fold or greater were considered significant. Gene ontology enrichment of differential genes was performed using Panther.

Polysome profiling of ovaries was adapted from previous protocols (McCarthy et al., 2022). Approximately 100 young wild type flies (*UAS-dcr;nosGAL4*) or about 275 experimental ovary pairs Nop60B were dissected (within 2 hrs of eclosion) in 1X PBS. The ovaries were immediately flash frozen on liquid nitrogen. Samples were homogenized by motorized pestle in lysis buffer and 20% of lysate was used as input for mRNA isolation and library preparation (as described above). Samples were loaded onto 10-45% CHX supplemented sucrose gradients in 9/16 x 3.5 PA tubes (Beckman Coulter, #331372) and spun at 35,000 x g in an SW41 rotor for 3 hours at 4°C. Gradients were fractionated with a Density Gradient Fractionation System (#621140007). RNA was extracted using acid phenol-chloroform and precipitated overnight. Pelleted RNA was resuspended in 20 μL water, treated with TURBO DNase and libraries were prepared as described above.

### Polysome-Seq Analysis

Analysis of polysome-seq was done using https://ruggleslab.shinyapps.io/RIVET/ (Ernlund et al., 2018). Polysome associated targets were further defined using the following parameters. Lowly associated mRNAs were identified by <-2 fold change and <0.05 p-Value while highly associated mRNAs were identified by >2 fold change and <0.05 p-Value.

### Ribosome footprinting

#### Ribo-Seq library preparation

Ribosome footprinting was performed as previously described (Dunn et al., 2013) with several modifications. ∼500 µL of ovaries were hand-dissected in Schneider’s Drosophila Medium (ThermoFisher), washed twice in 1 mL of lysis buffer (0.5% Triton X-100, 150 mM NaCl, 5 mM MgCl2, 50 mM Tris-HCl pH 7.5), and flash frozen in 2 mL of lysis buffer supplemented by 1 mM DTT, 50 µM GMP-PNP, 2 µg/mL emetine, and 20 U/mL Superase·In RNase Inhibitor (Ambion) in liquid N2. Ovaries were lysed using a Cellcrusher tissue pulverizer (Cellcrusher), allowed to thaw on ice, and centrifuged first at 10,000 rpm for 10 min and then at 13,200 rpm for 10 min. 300 µL of supernatant was used for footprint library preparation, and another 300 µL were used for poly(A)-selected mRNA-Seq library preparation. Ribosome footprints were generated by incubating the lysate with 3 U/µg of micrococcal nuclease (NEB) for 40 min at 25° C, then quenching by addition of EGTA to a final concentration of 6.25 mM. Ribosomes were sedimented through a 34% sucrose cushion for 2.5 hr at 33,000 rpm in a Beckman SW50 rotor, and the pellet was re-suspended in 10 mM Tris pH 7.0. RNA was extracted using TRIzol LS (Invitrogen) and size-selected (28-34 nt) on a 15% TBE-urea gel. RNA was then de-phosphorylated by incubating with T4 polynucleotide kinase (NEB) for 1 hr at 37° C, size-selected, and ligated to the 3′ adapter by incubating with T4 RNA ligase 2 truncated mutant (NEB) and 1 µg of pre-adenylated adapter (5′rAppCTGTAGGCACCATCAAT/3ddc) for 2 hr at 25° C. The ligation products were size-selected on a 10% TBE-urea gel. Reverse transcription was performed with Superscript III (Invitrogen) using the Illumina Tru-Seq RT primer:

/5Phos/AGATCGGAAGAGCGTCGTGTAGGGAAAGAGTGTAGATCTCGGTGGTCGC

/iSp18/CACTCA/iSp18/TTCAGACGTGTGCTCTTCCGATCTATTGATGGTGCCTACAG and the reaction was quenched by incubating with 0.1M NaOH for 20 min at 98° C. Following rRNA depletion, cDNA libraries were circularized by two sequential CircLigase (Epicentre) reactions and amplified by 9-12 PCR cycles.

#### mRNA-Seq library preparation

Total RNA was extracted from 300 µL of lysate with TRIzol LS, precipitated with isopropanol, washed in ice-cold 80% ethanol, and re-suspended in 10 mM Tris-HCl pH 7.0. mRNA-Seq libraries were then prepared from poly(A)-selected mRNA according to manufacturer’s instructions using the Illumina TruSeq RNA Library Prep Kit.

#### Processing of sequencing data

All steps were performed on the Princeton Galaxy server (galaxy.princeton.edu). Multiplexed libraries were de-multiplexed using the Barcode Splitter tool with up to 2 mismatches. Illumina Tru-Seq adapters were clipped using the Trim Galore! tool. The trimmed reads were first mapped against Drosophila rRNA sequences using Bowtie with default parameters, and the un-aligned reads were then aligned to the Drosophila genome Release 6 (dm6) using Bowtie2 with default parameters. The resulting BAM files were used for subsequent analyses.

#### Peak detection

The *Drosophila melanogaster* genome (dm6) was divided into 30-bp tiles and the number of reads aligned to each tile was reported using the bamCoverage tool of the deepTools 2 programming suite (Ramírez et al., 2016). Resulting bedgraph files were pre-processed to break up 30-bp tiles into 30 1-bp tiles (Script1). Peak detection was then performed in R using the Bioconductor software suite (Gentleman et al., 2004; Huber et al., 2015). Tiles were first aligned to the transcript regions by gene using the TxDb.Dmelanogaster.UCSC.dm6.ensGene annotation, rtracklayer (Lawrence et al., 2009), GenomicRanges (Lawrence et al., 2013), and BioPhysConnectoR (Hoffgaard et al., 2010) R packages (Script2). Then the distribution of coverage in the tiles aligned to each gene transcript region was fit to a normal distribution using the MASS R package (Venables et al., 2002) (Script3 and Function1). Finally, the coverage distribution and tiles aligned to each gene region were used to identify peak containing tiles (Script4 and Function2). Peak tiles from different ribosome profiling libraries were then compared (Function3) and the names, locations, and actual sequences of high confidence peaks were extracted (Script5, Script6, and Function4) using the Bsgenome.Dmelanogaster.UCSU.dm6 annotation and the Biostrings (H. Pagès, 2017) and GenomicRanges (Lawrence et al., 2013) R packages. Peaks present in at least two of the three Ribo-Seq libraries but not in the control RNA-Seq libraries at the corresponding positions were considered high confidence ribosome footprint peaks.

### Mass spectrometry

Ovaries were dissected in 1X PBS and homogenized by motorized pestle in 100uL of TRIzol (Invitrogen, 15596026). RNA was isolated by adding an additional 950 uL of TRIzol and 200uL of Chloroform with mixing. Samples were centrifuged at 13,000 rpm, 4°C for 15 minutes. The aqueous phase was transferred to a new tube. Nucleic acids were precipitated by adding and equal volume of 5 M Ammonium Acetate (Sigma-Aldrich), 2.5 volumes 100% ethanol and precipitated for >1 hour at −80°C. Samples were centrifuged at 13,000 rpm, 4°C for 20 minutes. Ethanol was decanted, pellets were washed four times with 1 mL of cold 70% ethanol and dried at room temperature for 10 minutes. Pellets were dissolved in 20 μL RNase free water and placed in a 42°C for 10 minutes.

RNA concentration was determined by using UV 260 nm. The RNA was then treated with nuclease P1 and phosphodiesterase to obtain the desired ribonucleotide monophosphate mixtures for mass spectrometric analysis, as previously described (McIntyre et al., 2018; Rose et al., 2016, 2015).

Immediately before analysis, the obtained mononucleotide mixtures were diluted to 4 ng/μL in 10 mM ammonium acetate and 10% isopropanol. All samples were analyzed on a Thermo Scientific LTQ-Orbitrap Velos instrument as previously described (Rose et al., 2016, 2015; Mclntyre et al., 2018). Analyses were accomplished by using direct infusion electrospray ionization (ESI) in negative ion mode.

The relative abundance of each RNA PTM was expressed as Abundance versus Proxy (AvP), which was calculated according to the following equation: 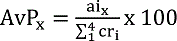 in which the signal intensity (ai_x_) of each RNA PTM was normalized against the sum of the intensities displayed in the same spectrum by the four canonical bases (cr_i_).

The RNA PTM profiling table translates relative abundances in AvP units to a hot-cold color gradient. The relative abundances displayed by the samples in the first column on the left were used as the baseline for comparisons with the rest of the samples. A different color was assigned only if the respective values were statistically different according to an unpaired student *t*-test with a *P*-value < 0.05. Each data point was the result of three to five biological replicates, which were each separately analyzed three times (technical replicates). Therefore, each value represents the average and standard deviation of a total of 9 to 15 separate analyses.

Tandem mass spectrometry was carried out in negative mode to differentiate uridine and pseudouridine (Rose et al., 2016, 2015; Mclntyre et al., 2018). The contribution of each isomer to the initial signal can be estimated from the relative intensities of their unique fragments. The abbreviations and complete names of each PTM in this study are available from MODOMICS (http://genesilico.pl/modomics/) database.

### Germline ribosome pulldowns

Ribosomal pulldowns were performed as previously described with the following modifications (Chen and Dickman, 2017). Approximately 50 young wild type ovaries (UAS-dcr;nosGAL4) and ∼100 Nop10^55194^ RNAi ovaries were dissected in PBS. After lysis in ribosomal lysis buffer, 120 uL was collected for input and Trizol extraction with previously described for mass spectrometry. The remaining lysate was divided into 180 uL aliquots. 6 ug of rabbit IgG (Jackson Immunoresearch) or rat-HA antibodies for 3 hours with rotation at 4°C. At hour 2, 50 uL Dynabeads A (Thermofisher) per replicate. The beads were prepped by performing four washes using a magnetic rack (500 uL for 2 minutes each) with ribosomal lysis buffer. After the fourth wash the beads were resuspended in 50 uL of ribosomal lysis buffer. To the samples either 25 uL of IgG or anti-HA was added and left overnight with rotation at 4°C. The following day, the beads were washed with 200 uL of ribosome lysis buffer for a total of four washes. After the final wash the beads were resuspended in 15uL of ribosome lysis buffer. A Trizol extraction was performed as previously described for mass spectrometry. After RNA extraction, a small portion of the RNA was run on a 1% agarose gel to confirm the presence of rRNA.

After the overnight incubation, the beads were washed with 200 uL of ribosome lysis buffer for a total of four washes. After the final wash the beads were resuspended in 15uL of ribosome lysis buffer. To the sample 4X SDS buffer was added and then heated at 95°C for 5 minutes and stored at −20°C until Western analysis.

### Western Blot

Ovaries were dissected in 1X PBS (Flora et al., 2018). After dissection, PBS was aspirated and 30 μl of RIPA buffer with protease inhibitors was added, and the tissue was homogenized. The homogenate was centrifuged at 13,000 rpm for 15 minutes at 4°C. The aqueous layer was transferred into a new tube while avoiding the top lipid layer. 1 μl of the protein extract was used to determine protein concentration via Invitrogen Qubit® Protein Assay Kit. 15-20 μg of protein was denatured with 4X Laemmli Sample Buffer and β-mercepthanol at 95°C for 5 minutes. The samples were loaded onto a Mini-PROTEAN TGX 4-20% gradient SDS-PAGE gels and run at 300V for 20 minutes. The proteins were then transferred to a 0.20 μm nitrocellulose membrane using Bio-Rad Trans-blot Turbo System. After transfer, the membrane was blocked in 5% milk in PBST for 2 hours at RT. The following antibodies were used: rat-HA (1:4000), rabbit-Vasa (1:6000) and rabbit-RpL26 (1:1000). Primary antibody was prepared in 5% milk in PBST was added to the membrane and incubated at 4°C overnight. The membrane was then washed three times in 0.5% milk PBST. Anti-rabbit HRP (1:10,000) or Anti-rabbit HRP (1:10,000) was prepared in 5% milk in PBST, and was added to the membrane and incubated at room temperature for 1 hour. The membrane was then washed 3 times in PBST. The Bio-Rad chemiluminescence ECL kit was used to image the membrane.

Note: To help normalize germline in the Western Blot probing for the PolyQ-HA reporter, first 15-20 ug of lysate run and probed for Vasa. Controls were then diluted 1:5 to help equalize the amount of germline loaded and compared to the H/ACA box member knockdown. Normalizations were performed using the top Vasa band.

### Stellaris *in situ* hybridizations

A modified in situ hybridization procedure for Drosophila ovaries was followed from (Sarkar et al., 2021). Probes were designed and generated by LGC Biosearch Technologies using Stellaris® RNA FISH Probe Designer, with specificity to target base pairs of target mRNAs. Ovaries (3 pairs per sample) were dissected in RNase free PBS and fixed as described above. The fixed tissue was washed with twice with 1 mL of PBS and then permabillized with 70% ethanol at 4°C for 2 hours. After permeabilzation, 1 mL of wash buffer was added (40 mL RNase free water, 5 mL deionized formamide and 5 ML 20X SSC) for a 5 minute wash. To the sample 50 uL of a Stellaris Hybridization buffer, 10% (vol.vol.) of formamide with 50-100 mm of oligos and properly diluted antibodies were added and incubated at 30°C for a minimum of 16 hours in the dark. After the overnight incubation, the sample was washed twice with 1 mL of wash buffer with properly diluted secondary antibodies for 30 minutes. After the second wash Vectashield was added and samples were imaged.

Stellaris probes were designed on (https://www.biosearchtech.com/support/education/stellaris-rna-fish) to all possible isoforms and Cy3 probe. Sequences found in excel file stellarisinsituprobes.xslx.

### Quantitative Real Time-PCR (qRT-PCR)

Once RNA was purified and isolated, see RNA Isolation section, a reverse transcription (RT) was performed using Superscript II according to the manufacture’s protocol with equivalent volumes of RNA for each sample. cDNA was amplified using 5μL of SYBR green Master Mix, 0.3 μL of 10μM of each reverse and forward primers in a 10 μL reaction. For each sample 3 biological and a minimum of 2 technical replicates were performed. Technical replicates were averaged, and tubulin was utilized as a control. To calculate fold change relative to tubulin mRNA levels, the average of the 2^-ΔCt for the biological replicates was calculated with error bars representing Standard Error of the Mean. Statistics were performed using a paired t-test on ΔCt values.

### MEME Analyses

The 5′UTR, CDS, 3′UTR and amino acid sequence of 465 mRNAs that are lowly associated with polysome than control and 320 mRNAs highly associated with polysome and analyzed by the MEME algorithm (Bailey, n.d.). Discriminative mode analysis was conducted against 1573 non-target gene sequences as background with default parameters. Motif logos, number of sites, and E-values all reported as produced by output of the program.

### FIMO Analyses

An amino acid motif of 5Qs was run against the amino acid sequences of all mRNAs that were lowly associated with polysome. Motifs identified in targets were searched in the given strand with a p-value < 1E-4.

---

**Supplemental 1:**
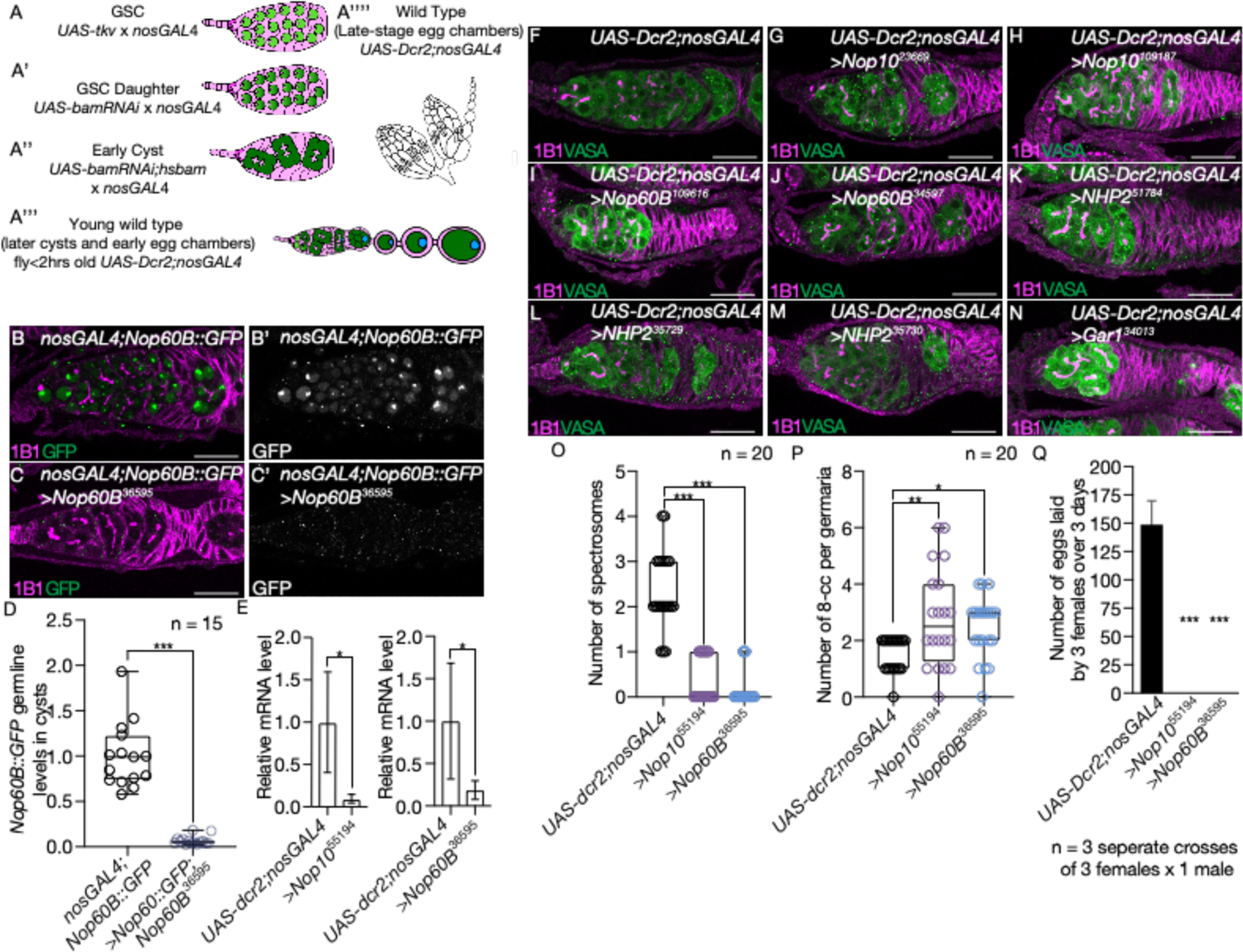
The H/ACA box is required in the germline for proper oogenesis (A-A’’’’) Schematic of method to enrich for GSCs (A), GSC daughters (A’), cysts (early cysts; A’’), young wild type (later cysts and early egg chambers; A’’’), and late-stage egg chambers (A”). (B,C) *nosGAL4;Nop60B::GFP* (driver control) germarium (B) and germarium with germline knockdown of *Nop60B* in Nop60B::GFP (C) stained with anti-1B1 (magenta) and anti-GFP (green). GFP is shown in gray scale (B’ and C’). Scale bar for all images is 20 µm. (D) Quantification using unpaired t-test of GFP in cysts of *nosGAL4;Nop60B::GFP* and germline knockdown of *Nop60B* in Nop60B::GFP background (n = 15 each, *** p<0.0001). There were lower levels of germline GFP in *Nop60B* knockdown germaria. (E) qRT-PCR assaying the RNA levels of *Nop10* or *Nop60B* in germline RNAi normalized to control, *UAS-Dcr;nosGAL4*, and indicating successful knockdown of H/ACA box members (n = 3, Nop10: * p = 0.0231, paired t-test) (n=3, Nop60: * p = 0.0142, paired t-test). Error bars representing SEM. (F-N) *UAS-Dcr2;nosGAL4* (driver control) germaria (F) and germline depletion of varying members of the H/ACA box stained with anti-1B1 (magenta) and anti-Vasa (green): *Nop10* (G-H); *Nop60B* (I-J); *NHP2* (K-M); and *Gar1* (N). Germline knockdown of H/ACA box members results in a block in cyst development. Scale bar for all images is 20 µm. (O) Quantification of number of spectrosomes in *UAS-Dcr2;nosGAL4* (driver control) germaria and germline depleted of *Nop10* or *Nop60B* showing a loss of GSCs. Statistics performed were Dunnett’s multiple comparisons test post-hoc test after one-way ANOVA (n = 20 each, *** p<0.0001). (P) Quantification of number of 8-cell cysts in *UAS-Dcr2;nosGAL4* (driver control) germaria and germline depleted of *Nop10* or *Nop60B* showing an increase in 8-cell cysts. Statistics performed were Dunnett’s multiple comparisons test post-hoc test after one-way ANOVA (n = 20 each, *p=0.0341, **p=0.0030,). (Q) Egg laying assay after germline RNAi knockdown of *Nop10* or *Nop60B* indicating a loss of fertility compared to *UAS-dcr2;nosGAL4* (driver control) (n = 0-173, *** p<0.001) Dunnett’s multiple comparisons test post-hoc test after one-way ANOVA, p < 0.0001. Error bars are standard deviation (SD).

**Supplemental 2:**
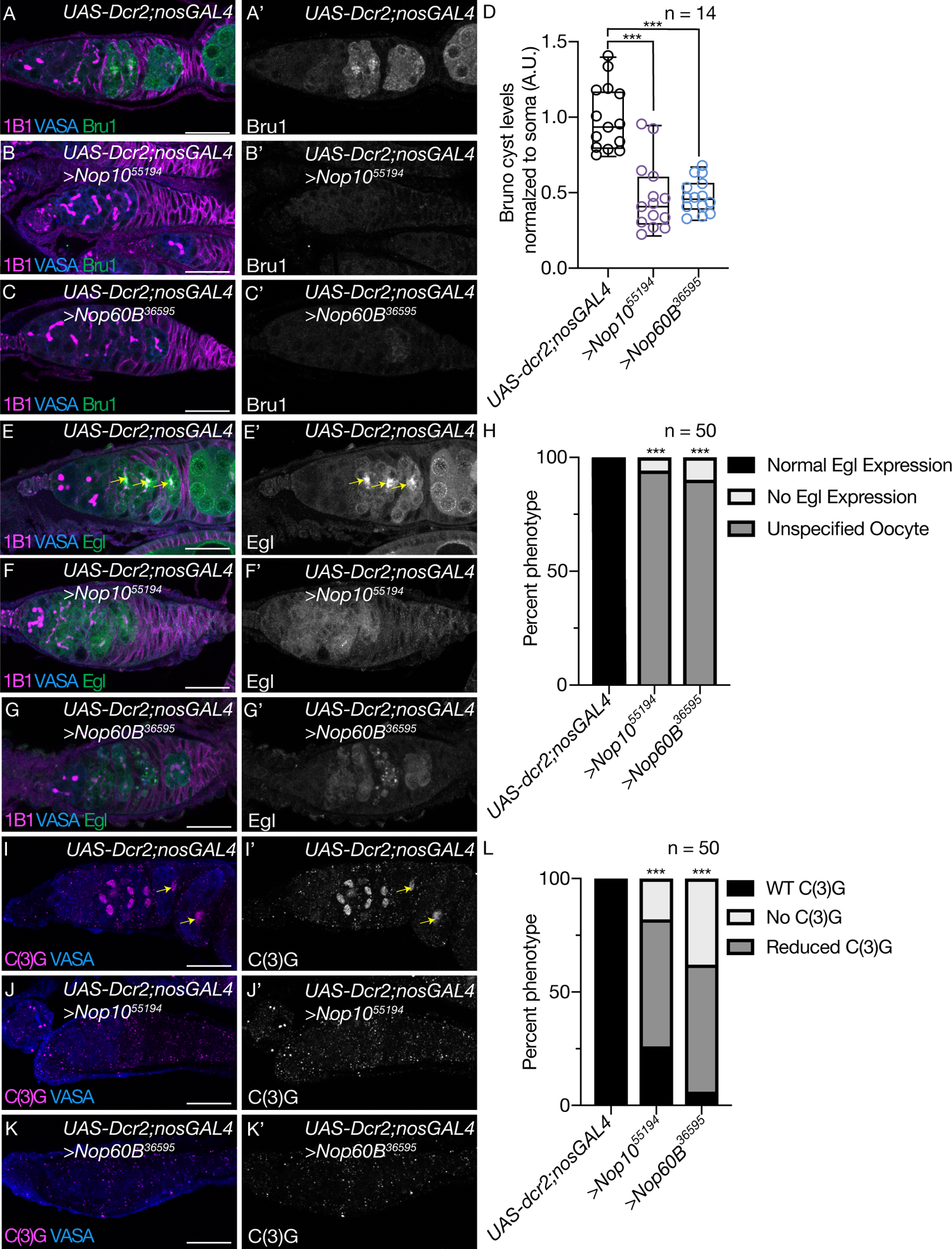
The H/ACA box is required for meiotic progression (A-C) *UAS-Dcr2;nosGAL4* control germaria (A) and germline depleted for *Nop10* and *Nop60B* (B, C) stained with anti-1B1 (magenta), anti-Vasa (blue) and anti-Bru1 (green). Bru1 is shown in gray scale (A’, B’, C’). (B) Quantification of Bru1 levels in germline depleted of *Nop10* and *Nop60B* normalized to soma showing a reduction in Bru1 levels in germaria depleted of the H/ACA box. Statistics performed were Dunnett’s multiple comparisons test post-hoc test after one-way ANOVA (n = 50 each, *** p<0.0001). (E) *UAS-Dcr2;nosGAL4* (driver control) germaria and (F, G) germline depleted of *Nop10* and *Nop60B*, stained with anti-1B1 (magenta), anti-Vasa (blue) and anti-Egl (green). Egl is shown in gray scale (E’, F’, G’). Arrow pointing at designated oocyte. Scale bar for all images is 20 µm. (H) Quantification of oogenesis defect phenotypes showing a loss of oocyte specification in germaria depleted of the H/ACA box. Statistical analysis performed with Fisher’s exact test (n = 50 each, *** p<0.0001). (I-K) *UAS-Dcr2;nosGAL4* (driver control) germaria (I) and germline depleted of *Nop10* (J) or *Nop60B* (K) stained with anti-c(3)G (magenta) and anti-Vasa (blue). c(3)G is shown in gray scale (I’, J’, K’). Arrow points to designated oocyte. Scale bar for all images is 20 µm. (L) Quantification of oogenesis defect phenotypes showing a loss of c(3)G expression in germaria depleted of the H/ACA box. Statistical analysis performed with Fisher’s exact test (n = 50 each, *** p<0.0001).

**Supplemental 3:**
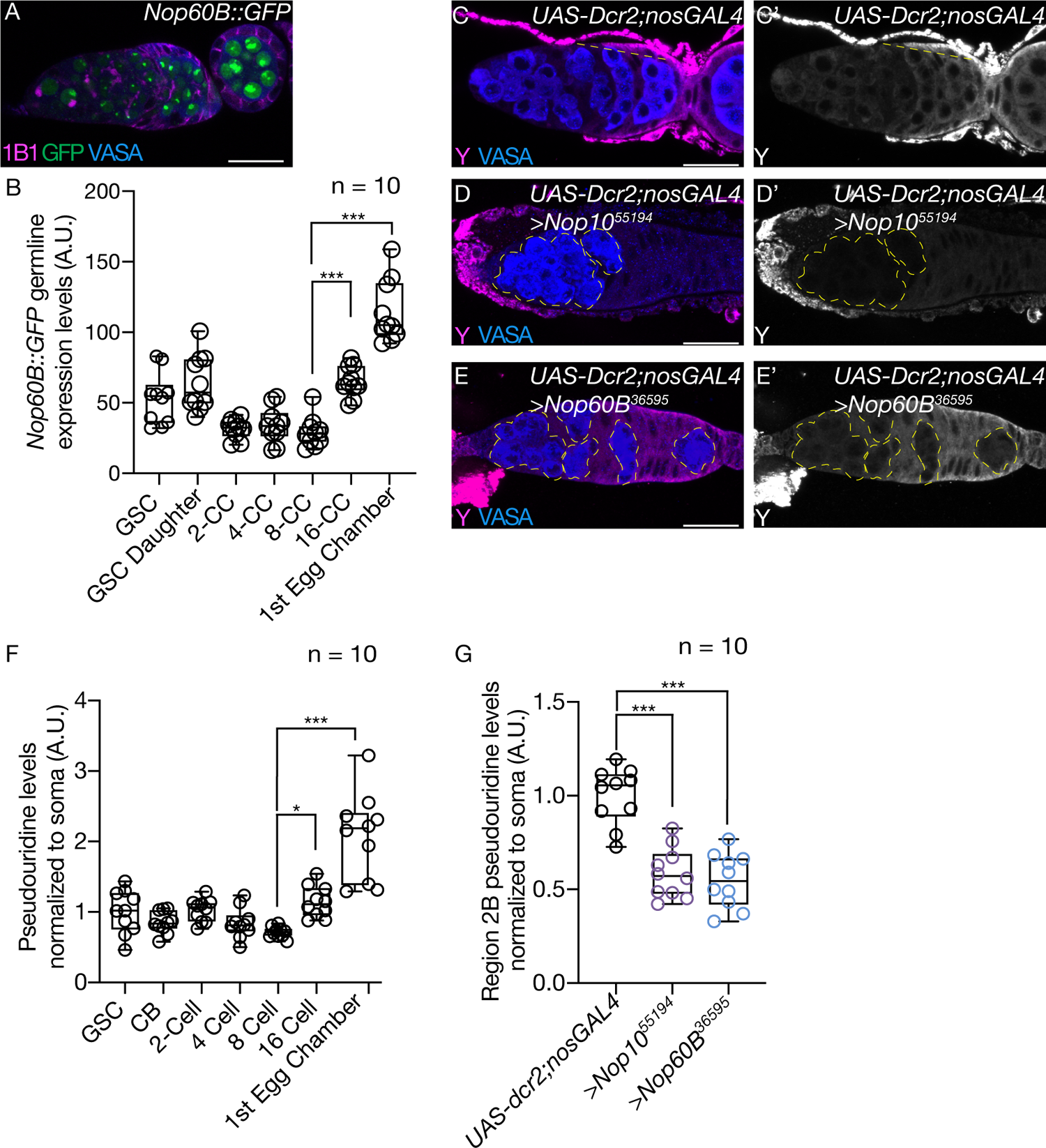
Nop60B and pseudouridine increase during transition from cyst to egg chamber (A) *Nop60B::GFP* germarium stained with anti-1B1 (magenta), anti-Vasa (blue) and anti-GFP (green). GFP is shown in gray scale (A’). Image is taken at 40x and scale bar for all images is 20 µm. (B) Quantitation of GFP from GSC to 1^st^ egg chamber in *Nop60B::GFP*. Statistics performed were Tukey’s multiple comparisons test post-hoc test after one-way ANOVA. Statistics shown comparing 8-cell cyst to 16-cell cyst and to the egg chamber showing an increase in GFP levels (n = 10, *** p<0.001). (C) *UAS-Dcr2;nosGAL4* (driver control) germaria and (D, E) germline depletion of *Nop10* and *Nop60B* stained with anti-pseudouridine (magenta) and anti-Vasa (blue). Pseudouridine is shown in gray scale (C’, D’ and E’). Yellow dotted line in control represents area of increasing pseudouridine levels while yellow outline in *Nop10* and *Nop60B* represents loss of pseudouiridine. Scale bar for all images is 20 µm. (F) Quantification of pseudouridine levels from GSC to 1^st^ egg chamber in *UAS-Dcr2;nosGAL4*. Statistics performed were Tukey’s multiple comparisons test post-hoc test after one-way ANOVA. Statistics shown comparing 8-cell cyst to 16-cell cyst and to the egg chamber showing an increase in pseudouridine levels (n = 10, * P < 0.0313, *** p< 0.001). (G) Statistics shown from the 2B-region of *UAS-Dcr2;nosGAL4* and *Nop10* or *Nop60* depleted germaria. Loss of H/ACA box members led to a significant reduction in germline pseudouridine levels when normalized to soma (n = 10, *** p< 0.001).

**Supplemental 4:**
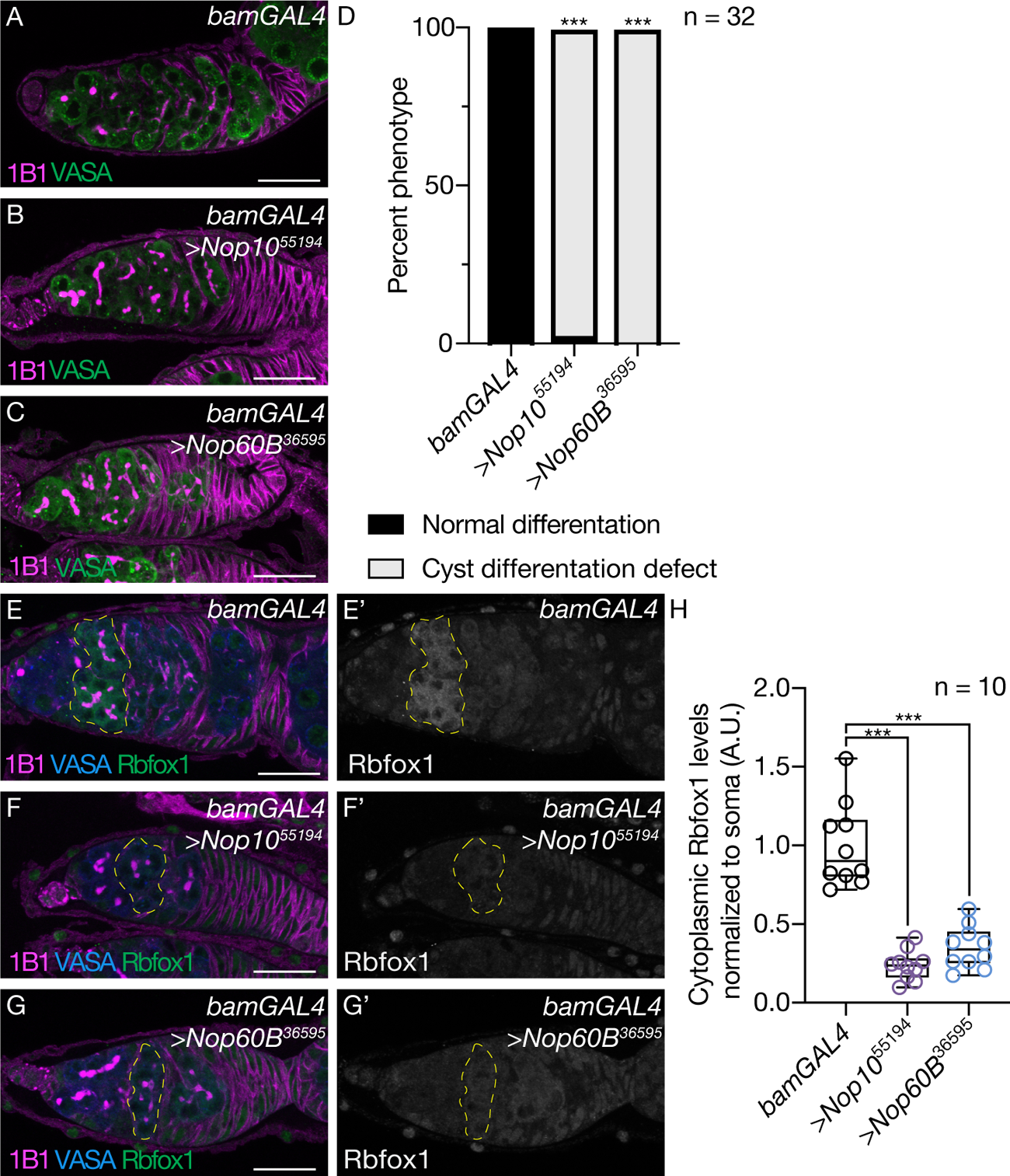
The H/ACA box is required in the cyst stages (A-C) *bamGAL4* (driver control) germaria (A) and germline depletion of the *Nop10* (B) or *Nop60B* (C) stained with anti-1B1 (magenta) and anti-Vasa (green). Scale bar for all images is 20 µm. (D) Quantification of oogenesis defect phenotypes per genotype showing a cyst differentiation defect. Statistical analysis performed with Fisher’s exact test (n = 32 for all, *** p<0.0001). (E) *bamGAL4* (driver control) germaria and (F and G) germline depletion of *Nop10* and *Nop60B* stained with anti-1B1 (magenta), anti-Vasa (blue) and anti-Rbfox1 (green). Rbfox1 is shown in gray scale (E’, F’ and G’). Scale bar for all images is 20 µm for all images. Yellow dotted line outlined cysts. (H) Quantification of Rbfox1 levels showing a reduction in Rbfox1 levels in *Nop10* and *Nop60B* depleted germaria. Statistical analysis performed with Fisher’s exact test (n = 10 each, *** p<0.0001).

**Supplemental 5:**
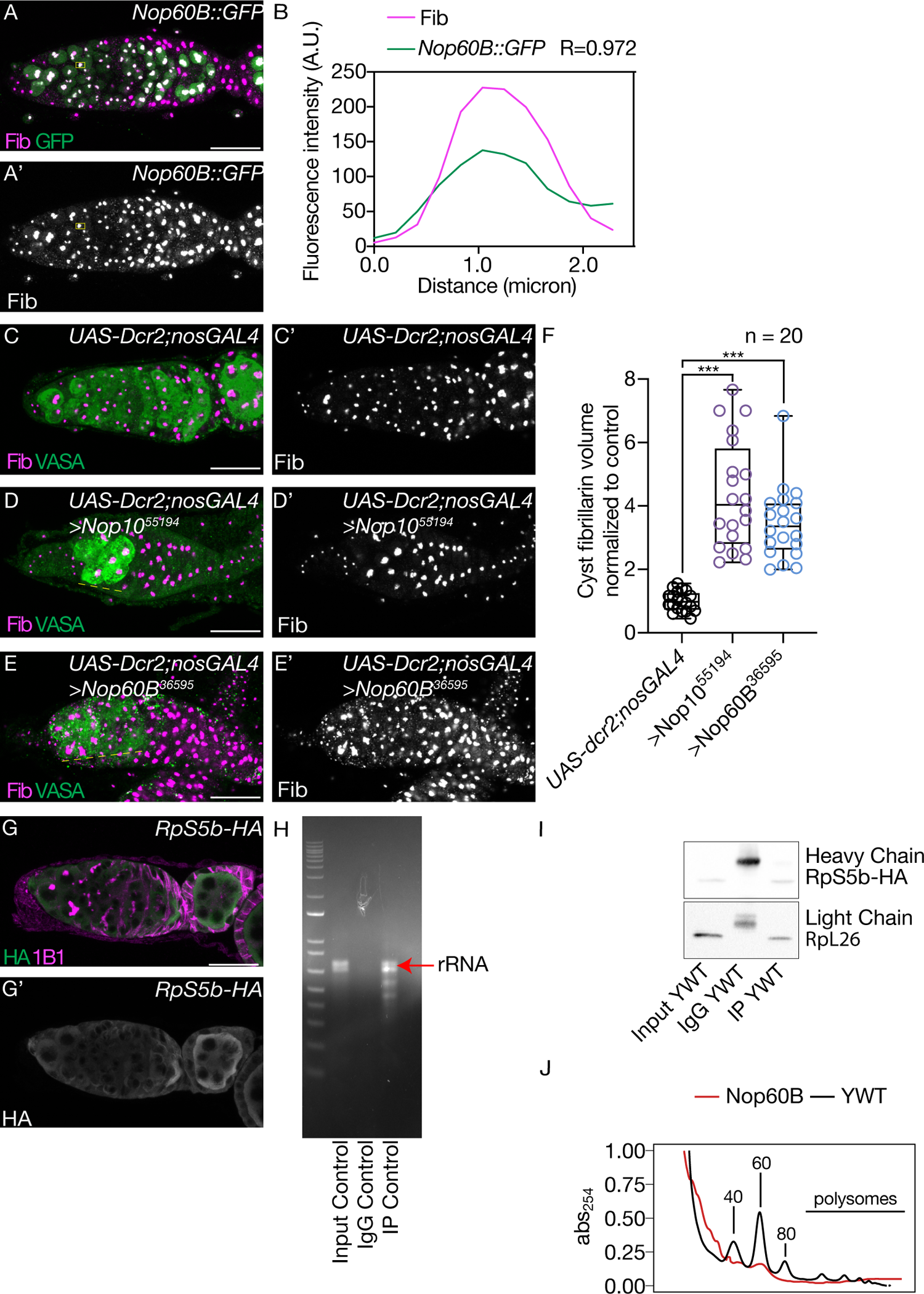
The H/ACA box deposits pseuoduiridne on rRNA and is required for proper ribosome biogenesis (A) *Nop60B::GFP* germarium stained with Fibrillarin (magenta) and GFP (green). Fibrillarin is shown in gray scale (A’). Scale bar for all images is 20 µm. (B) Fluorescence intensity plot generated from a box of averaged pixels centered around the punctate of Fibrillarin in the white box. R values denote Spearman correlation coefficients between GFP and Fibrillarin from plot profiles generated using Fiji, taken from the nucleolus denoted by the yellow box. (C) *UAS-Dcr2;nosGAL4* (driver control) germaria and (D and E) germline depletion of *Nop10* and *Nop60B* stained with fibrillarin (magenta) and Vasa (green). Fibrillarin is shown in gray scale (C’, D’ and E’). Scale bar for all images is 20 µm. (F) Quantification of nucleolar volume in the cysts stages per genotype showing an increased nucleolar size with *Nop10* and *Nop60B* depletion when compared to control. Statistics performed were Dunnett’s multiple comparisons test post-hoc test after one-way ANOVA (n = 20 each, *** p<0.0001). (G) Germaria of *RpS5b-HA* stained with anti-1B1 (magenta) and anti-HA (green). HA is shown in gray scale (G’). (H) Agarose gel of control lysate showing enrichment of rRNA (red arrow) in the input and IP lane but not in negative control (IgG). (I) Western blot analysis of ribosomal pulldowns probing for HA and RpL26 in input, IgG and pulldown samples. HA and RpL26 are present in both the input and pulldown lane but not the IgG showing successful pulldown of large and small ribosomal proteins. (J) Polysomes traces for YWT (*UAS-Dcr2;nosGAL4)* (black) and *Nop60B* (red) depleted germaria. Nop60B is required for proper ribosome biogenesis as traces show that loss of *Nop60B* results in 40S and 60S defects as well as loss of polysomes when compared to control.

**Supplemental 6:**
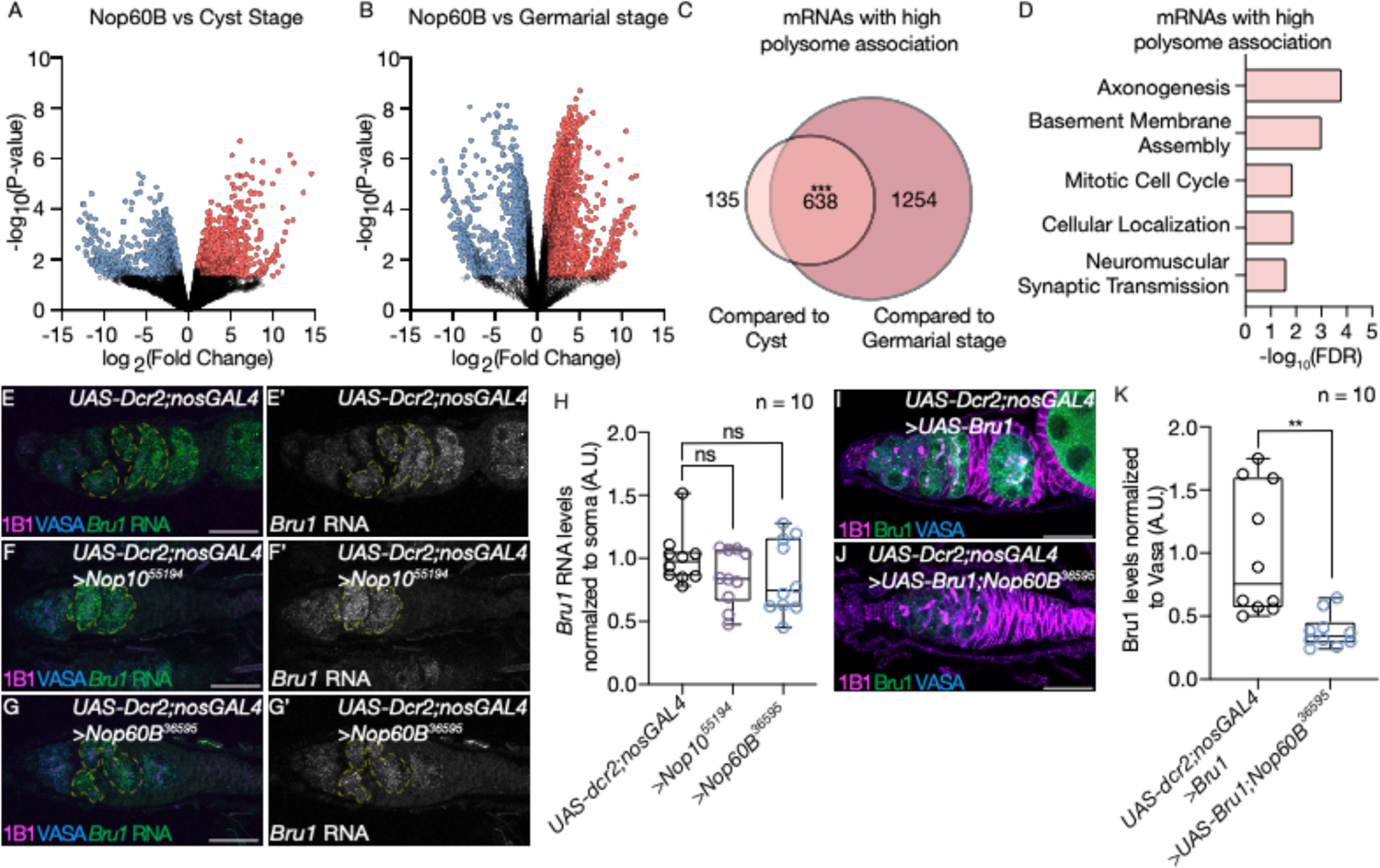
The H/ACA box is required for translation of meiotic mRNAs (A) Volcano plot of *Nop60B* depleted germaria vs cyst stages (heat shock) with log_2_(fold change) on x-axis and –log_10_(P-value) on the y-axis. Blue points represent mRNAs that have lower association with the polysomes and red points represent mRNAs with high polysome association (n = 2, targets identified as 2-fold cutoff). (B) Volcano plot of *Nop60B* depleted germaria vs germarial stages (YWT or *UAS-Dcr2;nosGAL4*) with log_2_(fold change) on x-axis and –log_10_(P-value) on the y-axis. Blue points represent mRNAs that have lower association with the polysomes and red points represent mRNAs with high polysome association (n = 2, targets identified as 2-fold cutoff). (C) Venn diagram illustrating overlap of Nop60B-polysome >2 fold more association upon loss of *Nop60B* (significance to low to compute in RStudio using Hypergeometric Test). Controls were cysts, enriched through heat shock, and germarial stages (YWT or *UAS-Dcr2;nosGAL4*) (D) Significant biological process GO terms of shared highly associated mRNAs in ovaries depleted of *Nop60B* compared to control sets, showing an enrichment for mRNAs associated with mitotic cell cycle. (E-G) In situ hybridization to *Bru1* RNA (green) together with anti-1B1 (magenta) and anti-Vasa (blue) staining in *UAS-Dcr2;nosGAL4* (driver control) germaria (E) and germline depleted of *Nop10* (F) and *Nop60B* (G). *Bru1* RNA is shown in gray scale (E’, F’, G’). Scale bar for all images is 20 µm. Yellow dotted line outlines *bru1* RNA. (H) Quantification of *Bru1* RNA levels in germline depleted of varying members of *Nop10* and *Nop60B* normalized to soma showing no significant change in *Bru1* RNA levels. Statistics performed were Dunnett’s multiple comparisons test post-hoc test after one-way ANOVA (n = 10 each, not significant, p=0.3606 and p=0.3752 respectively). (I) Germaria of *UAS-Dcr2;nosGAL4* (driver control) overexpressing Bru1 and (J) germline depleted of *Nop60B* overexpressing Bru1. Scale bar for all images is 20 µm. (K) Quantification of Bru1 levels in control vs germline depleted of *Nop60B* normalized to Vasa show a reduction in Bru1 levels. Statistics performed were unpaired t-test (n = 10 each, ** p=.0015).

**Supplemental 7:**
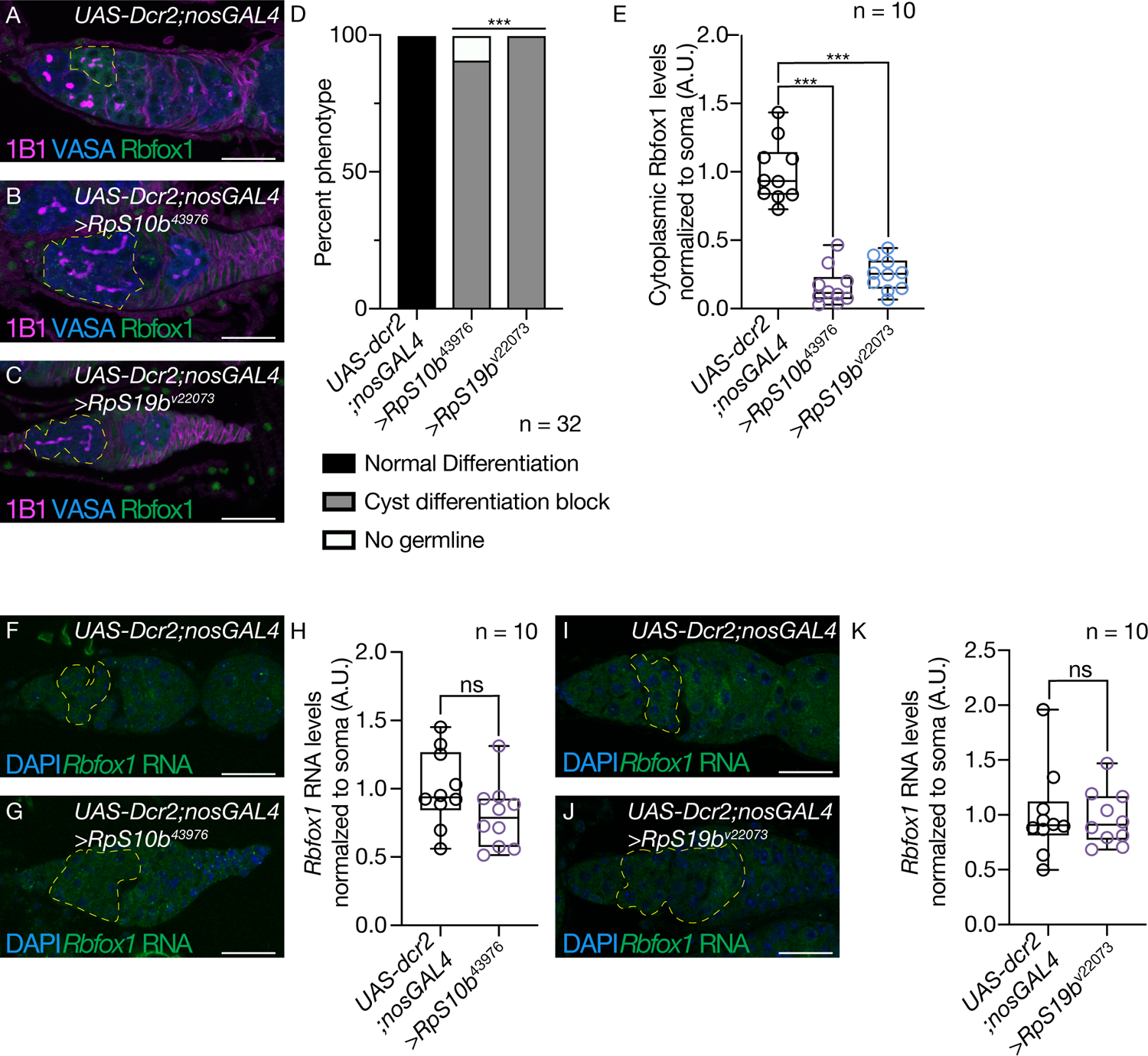
Ribosomal paralogs are required for Rbfox1 translation (A-C) *UAS-Dcr2;nosGAL4* (driver control) germaria (B) germline depletion of *RpS10b* (B) or *RpS19b* (C) stained with anti-1B1 (magenta), anti-Vasa (blue) and anti-Rbfox1 (green). Scale bar for all images is 20 µm. Yellow dotted lines outline cysts. (D) Quantification of oogenesis defect phenotypes per genotype. Knockdown of ribosomal paralogs results in a cyst differentiation block. Statistical analysis performed with Fisher’s exact test (n = 32 for all, *** p<0.0001). (E) Quantification of cytoplasmic Rbfox1 levels normalized to soma in germline depletion of *RpS10b* and *RpS19b* showing that loss of ribosomal proteins results in lower Rbfox1 levels. Statistics performed were Dunnett’s multiple comparisons test post-hoc test after one-way ANOVA (n = 10 each, *** p<0.0001). (F) In situ hybridization of *Rbfox1* RNA (green) and DAPI (blue) in *UAS-Dcr2;nosGAL4* (driver control) germaria and (G) germline depleted of *RpS10b*. Scale bar for all images is 20 µm. Yellow dotted line outlines *Rbfox1* RNA. (H) Quantification of *Rbfox1* RNA levels in germline depleted of *RpS10b* normalized to soma showing no significant differences in *Rbfox1* RNA levels. Statistics performed were unpaired t-test (n = 10 each, not significant, p=0.1006). (I) In situ hybridization of *Rbfox1* RNA (green) and DAPI staining (blue) in *UAS-Dcr2;nosGAL4* (driver control) germaria and (J) germline depleted of *RpS19b*. Scale bar for all images is 20 µm. Yellow dotted line outlines *Rbfox1* RNA. (K) Quantification of *Rbfox1* RNA levels in germline depleted of *RpS19b* normalized to soma showing no significant differences in *Rbfox1* RNA levels. Statistics performed were unpaired t-test (n = 10 each, not significant, p=0.8258).

**Supplemental 8:**
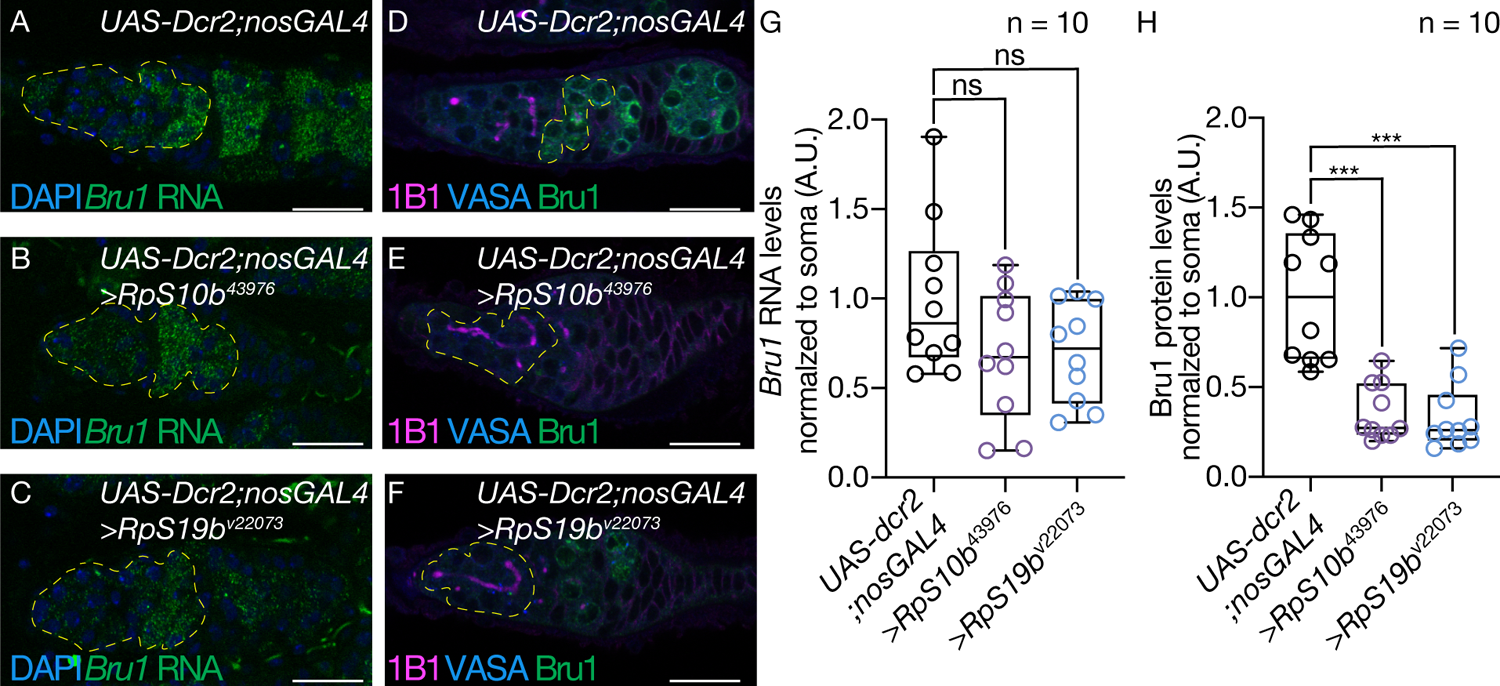
Ribosomal paralogs are required for Bru1 translation (A-C) In situ hybridization to *Bru1* RNA (green) and DAPI staining (blue) in *UAS-Dcr2;nosGAL4* (driver control) germaria (A) and germline depleted of *RpS10b* (B) and *RpS19b* (C). Scale bar for all images is 20 µm. Yellow dotted line outlines *bru1* RNA. (D-F) *UAS-Dcr2;nosGAL4* (driver control) germaria (D) and germline depletion of *RpS10b* (E) and *RpS19b* (F) stained with anti-1B1 (magenta), anti-Vasa (blue) and anti-Bru1 (green). Scale bar for all images is 20 µm. Yellow dotted line outlines cysts. (G) Quantification of *Bru1* RNA levels normalized to soma in germline depletion of *RpS10b* and *RpS19b* showing no significant differences in *Bru1* RNA levels with loss of ribosomal paralogs. Statistics performed were Dunnett’s multiple comparisons test post-hoc test after one-way ANOVA (n = 10 each, not significant, p=0.1149 and 0.1325, respectively). (H) Quantification of Bru1 protein levels normalized to soma in germline depletion of *RpS10b* and *RpS19b* showing that loss of ribosomal proteins results in lower bru1 levels. Statistics performed were Dunnett’s multiple comparisons test post-hoc test after one-way ANOVA (n = 10 each, *** p<0.0001).

**Supplemental 9:**
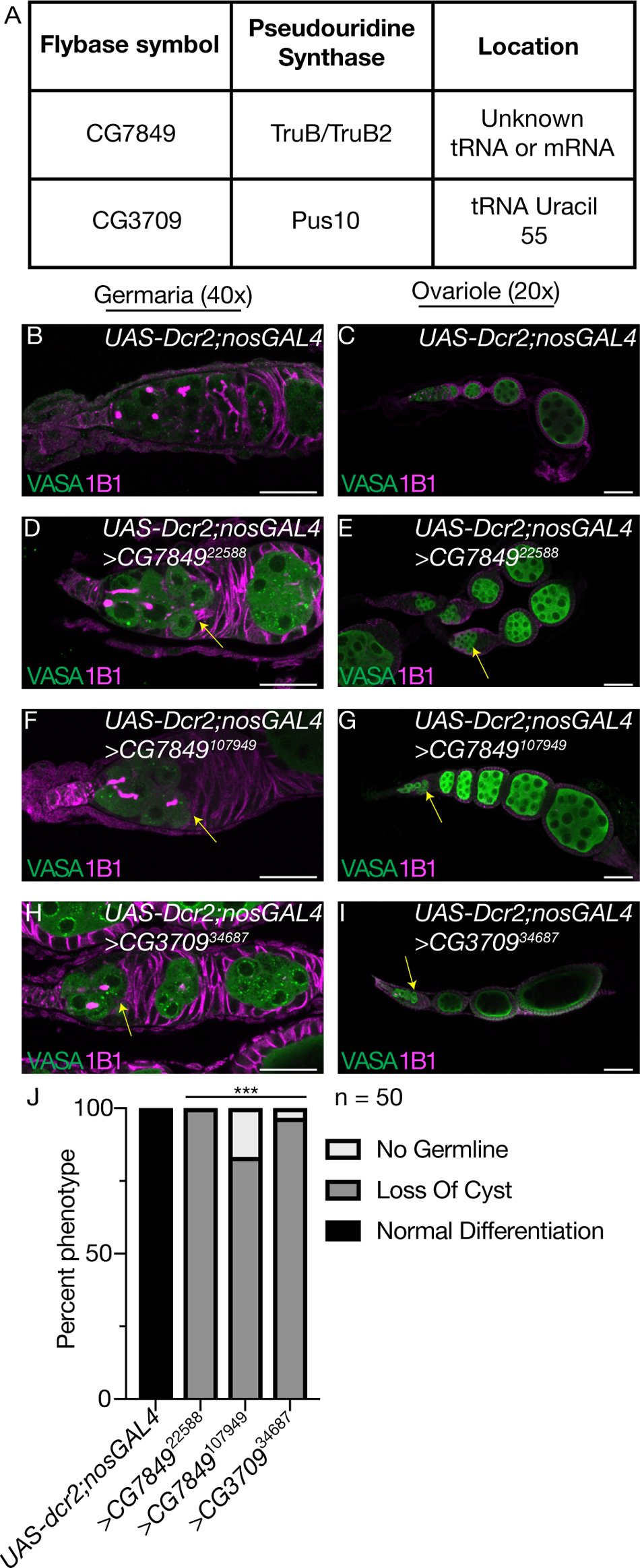
tRNA pseudouridine synthases are required for differentiation but do not phenocopy loss of rRNA pseudouridine synthases. (A) Table of tRNA pseudouridine synthases and location of pseudouridine deposition found to have a differentiation defect. (B, C) Images of 40x *UAS-Dcr2;nosGAL4* (driver control) germarium (B) and 20x *UAS-dcr2;nosGAL4* (driver control) ovarioles (C) stained with anti-1B1 (magenta) and anti-Vasa (green). (D, E) Images at 40x (D) and 20x (E) of germarium where *CG7849* is depleted in the germline and stained with anti-1B1 (magenta) and anti-Vasa (green). (F, G) Images at 40x (F) and 20x (G) of germarium using a second RNAi line to deplete *CG7849* in the germline and stained with anti-1B1 (magenta) and anti-Vasa (green). (H, I) Images at 40x (H) and 20x (I) of germarium where *CG3709* is depleted in the germline and stained with anti-1B1 (magenta) and anti-Vasa (green). Yellow arrow points to region where cysts are lost in all 20x images. Scale bar for all images is 20 µm. (J) Quantification of oogenesis defect phenotypes in tRNA pseudouridine synthase germline knockdowns resulting in loss of cyst defect. Statistical analysis performed with Fisher’s exact test (n = 50 each, *** p<0.0001).

**Supplemental 10:**
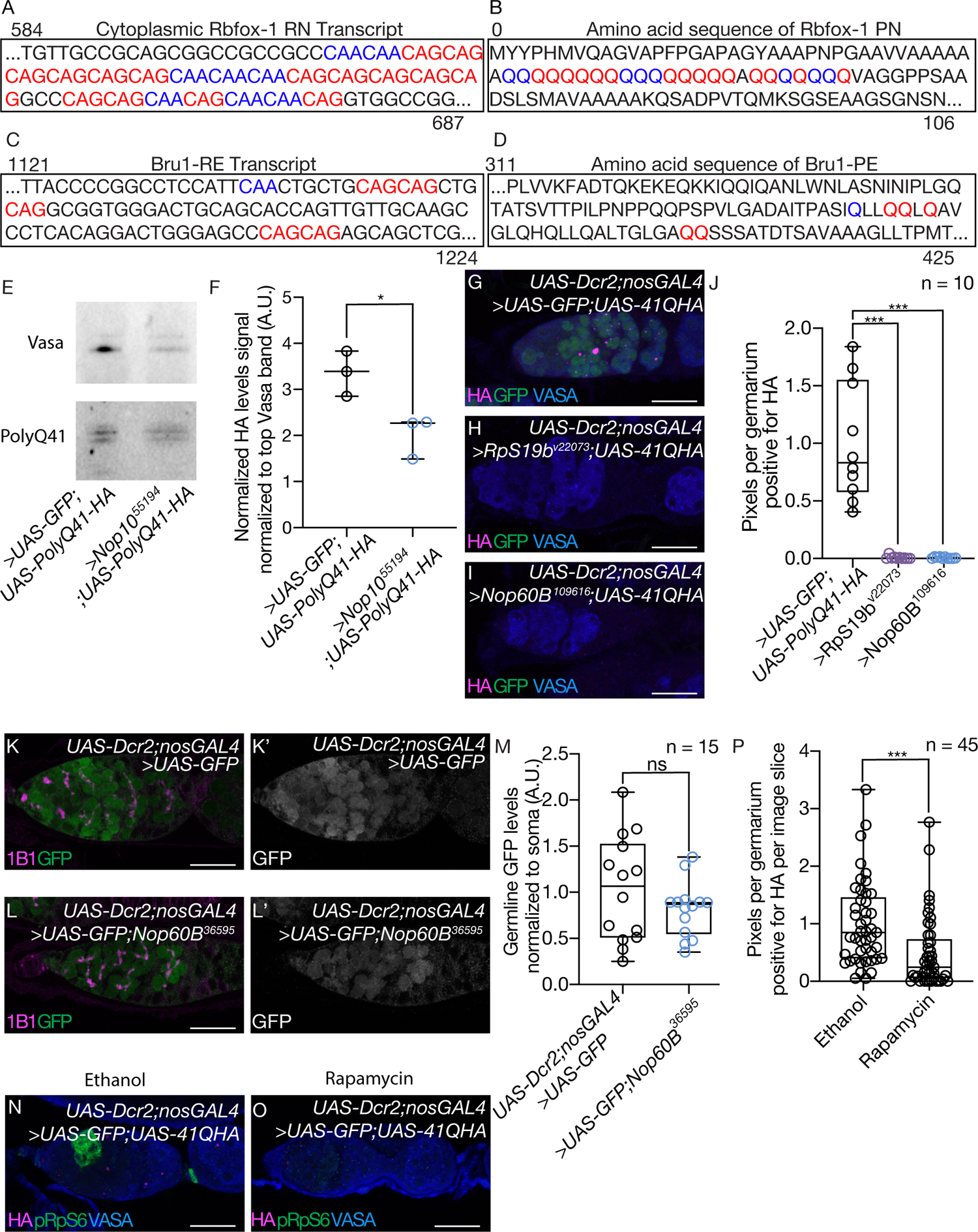
The H/ACA box is required for translating polyQ proteins (A) mRNA sequence of cytoplasmic Rbfox1-RN with glutamine (Q) codons CAA in blue and CAG in red. The transcript contains a large region of repeating CAA and CAG. (B) Protein sequence of Rbfox1-PN. Blue letters represent a Q encoded by CAA while red letters represent Q encoded by CAG. The transcript contains a large polyQ region. (C) mRNA sequence of Bru1-RE with CAA in red and CAG in blue. The transcript contains a large region of repeating CAA and CAG. (D) Protein sequence of Bru1-PE with Q encoded by CAA in blue, while red letter represent Q that corresponds to the codon CAG. The transcript contains a large polyQ region Q. (E) Western blot analysis of poly41Q-HA reporter driven in control and *Nop10* depleted germaria driven by *UAS-Dcr2;nosGAL4*. Western was probed with HA to detect polyQ protein. Vasa was probed for normalization of germline. (F) The level of HA (polyQ-HA) in ovary are significantly reduced upon germline knockdown of *Nop10*. Protein immunoblots for HA were performed using extracts from whole ovaries. The signal ratio between the HA and the upper Vasa band were used to quantitate and normalize the amount of germline. The ratio is expressed in arbitrary units (A.U.). The results of each independent experiment are plotted. Statistics performed were unpaired t-test (n = 3, * p=.0253). (G) Control confocal image of poly41Q-HA reporter driven in *UAS-Dcr2;nosGAL4* and germaria depleted of *RpS19b* and *Nop60B* (H and I) stained with anti-HA (magenta), anti-GFP (green) and anti-Vasa (blue). Scale bar for all images is 20 µm. (J) Quantification of percent of pixels per area of HA in control vs germline depleted of *RpS19b* and *Nop60B* showing a reduction in HA signal. Statistics performed were unpaired t-test (n = 10 each, *** p=0.0001). (K, L) UAS-GFP driven by *UAS-Dcr2;nosGAL4* in control germaria (K) and in germaria depleted of *Nop60B* (L), stained with anti-1B1 (magenta) and anti-GFP (green). GFP is shown in gray scale (K’ and L’). Scale bar for all images is 20 µm. (M) Quantitation of GFP levels in the cysts stages normalized to somatic background per genotype. There is no significant difference in GFP levels between control germaria and *Nop60B* depleted germaria. The results of each independent experiment are plotted. Statistics performed were unpaired t-test (n = 15, not significant, p=0.2187). (N, O) Confocal image of poly41Q-HA reporter driven by *UAS-Dcr2;nosGAL4* in mock-treated (N) and rapamycin-treated (O) ovaries, stained with anti-HA (magenta), anti-pRpS6 (green) and anti-Vasa (blue). Scale bar for all images is 20 µm. (P) Quantification of percent of pixels of HA per area in mock- and rapamycin-treated flies showing a reduction in HA signal with rapamycin treatment. Statistics performed were unpaired t-test (n = 45 slices quantified for each, *** p<0.0005).

**Table 1:** (A) PTM code for the modifications identified. The 1^st^ column represents the Modomics code, the 2^nd^ column represents the PTM name and the 3^rd^ column the shortened modification name. (B) Summary of RNA PTMs profiles obtained from GSCs, GSC daughters, cysts (early cysts), young wild type (later cysts and early egg chambers) and wild type (late-stage egg chambers). Each value represents the average and standard deviation of the respective relative abundances (AvP%, see Methods). A different shade of color was assigned only if the RNA PTMs relative abundance was statistically different from that of the GSCs input reference (1st column) with a p value not exceeding 0.05.

**Table 2:** (A) Excel spreadsheet of the RNA modification screen that contains the gene names, stock numbers, type of modification and phenotype. The raw number of germaria were counted. (B) The RNA modification screen represented as percent phenotypes.

**Table 3:** Summary of PTM profiles obtained. Each value represents the average and standard deviation of the respective relative abundances (AvP%, see Methods). A different shade of color was assigned only if the RNA PTMs relative abundance was statistically different from that of the cysts input reference (1st column) with a p value not exceeding 0.05.

**Table 4:** Spreadsheet of mRNA targets identified from pull-down utilizing pseudouridine antibody with a 2-fold cut off. (A) Genes that were lower than 2-fold enriched (B) genes that were higher thant 2-fold enriched and (C) fold-enrichment values for all genes.

**Table 5:** (A) MEME discriminative mode motif enrichment output of the 5’ UTR, CDS and 3’ UTR of genes that are lowly associated with the ribosome in germaria depleted of Nop60B. E-value, sites and width are provided for each identified motif. (B) MEME discriminative mode motif enrichment output of the 5’ UTR, CDS and 3’ UTR of genes highly associated polysome in germaria depleted of Nop60B depletion. E-value, sites and width are provided for each identified motif.

**Table 6:** (A) Correlation plots comparing Ribo-Seq datasets showing high reproducibility between libraries. (B) Column A: mRNA targets identified by Ribo-Seq that contain the CAG motif. Column B: mRNAs containing a strict repeating CAG (no interruptions). Column C: locations of the CAG motif. Column D: length of the longest CAG repeat present in the mRNA or if there are other amino acid repeats present.

**Table 7:** (A) Find Individual Motif Occurrences (FIMO) output of QQQQQ motif search in genes that were lowly associated with the polysome in Nop60B depleted germaria. Representative of 181 unique genes that significantly contain a motif resembling QQQQQ. (B) All transcripts from the Find Individual Motif Occurrences (FIMO) output of QQQQQ motif search in genes lowly associated with the polysome in Nop60B depleted germaria. Also provided are the p-value and matched motif sequences in each transcript.

